# Programmable synthetic cytokine receptors polarize macrophages to user-defined functional states

**DOI:** 10.64898/2026.05.12.724672

**Authors:** Judith C. Lunger, Lucas E. Sant’Anna, Antonio Salcido-Alcántar, Rebeca Arroyo Hornero, Wansang Cho, Alun Vaughan-Jackson, Mingxin Gu, Jenny Y. Liu, Alex N. Beckett, Joaquin Parrilla-Garcia, Sneha Ramakrishna, Michael C. Bassik, Kyle G. Daniels

## Abstract

Technology that precisely controls macrophage polarization to distinct functional states would deepen our understanding of macrophage biology and enable the development of new macrophage cell therapies. Here, we use a synthetic cytokine receptor (SCR) platform with a programmable signaling domain to control the polarization of primary human macrophages. SCRs containing signaling motifs from the interferon-gamma (IFN-γ) or Interleukin-10 (IL-10) receptors mimic key features of pro-inflammatory or anti-inflammatory polarization, respectively. Random recombination of nine distinct signaling motifs to create new SCR signaling domains generates a diverse landscape of synthetic macrophage states with varied expression of inflammatory markers (CD80, CD40) and anti-inflammatory markers (CD163, CD206), and varied phagocytic capacity. SCRs programmed with multiple YLxQ motifs increase macrophage phagocytosis of *E. coli* and chimeric antigen receptor (CAR)-macrophage phagocytosis of cancer cells in mice, reducing tumor burden by 30-fold. The motif-dependent polarization is well-described by a two-state model, enabling quantitative prediction of macrophage polarization state from SCR signaling domain composition. Leveraging this model, we design an SCR that simultaneously enhances phagocytosis and maintains a macrophage pro-inflammatory state. Together, these findings establish a framework for synthetic programming of macrophage polarization states, with potential applications in cancer immunotherapy and other disease contexts.

## Introduction

Macrophages are highly plastic cells that play critical roles in natural immunity and, increasingly, as engineered cell therapies^1^. Macrophages sense infection and tissue damage, phagocytose pathogens and debris, and sculpt the local environment through a broad repertoire of cytokines and chemokines^2^. Central to these roles is macrophages’ ability to polarize into distinct cell states with unique functional properties in response to external cues. For example, interferon-gamma (IFN-γ) can polarize macrophages into a state characterized by the display of costimulatory molecules and secretion of pro-inflammatory cytokines^3^. Conversely, interleukin (IL)-10 can polarize macrophages to a state characterized by the secretion of anti-inflammatory cytokines and promotion of tissue repair^4^. Evidence indicates that macrophages can integrate signals from multiple stimuli to adopt a broad spectrum of phenotypic states^5–7^. This plasticity enables macrophages to perform a variety of essential functions, such as secretion of pro-inflammatory or anti-inflammatory cytokines, clearance of apoptotic cells, and presentation of antigens. Dysregulation of such functions through aberrant polarization is associated with a range of diseases, including susceptibility to infection, cancer progression, atherosclerosis, and fibrosis^8–11^. Accordingly, there has been growing interest in engineering primary human macrophages for therapeutic applications. Technologies for precisely engineering macrophage polarization states would significantly enhance our understanding of macrophage biology and provide new strategies for controlling the immune landscape in various disease contexts.

Several strategies have advanced macrophage engineering, including polarization with exogenous cytokines, use of chimeric switch receptors^12^ or chimeric antigen receptors with polarizing signaling domains^13^, and gene knockdown or overexpression^14^. These approaches can guide macrophages to altered polarization states and enable coarse-grained control of macrophage phenotype. A remaining challenge is to reliably program macrophage polarization states for research and therapeutic applications. Ideally, we would be able to co-optimize phenotypes like phagocytic activity and inflammatory state. One potential solution is a programmable platform that allows the user to precisely navigate the polarization landscape and predictably guide human primary macrophages to desired states.

Cytokines are major drivers of macrophage polarization state^15^. In addition to IFN-γ and IL-10, cytokines like IL-4, IL-13, tumor necrosis factor (TNF), transforming growth factor (TGF)-β, macrophage colony-stimulating factor (M-CSF), granulocyte-macrophage colony-stimulating factor (GM-CSF), and others can alter macrophage polarization^6^. These cytokines are sensed by receptors whose intracellular domains contain short linear signaling motifs^16–18^. These signaling motifs serve as docking sites for downstream effectors that activate signaling cascades and drive phenotype changes^17,18^. Nature has evolved hundreds of cytokine receptors that use varied signaling motifs to steer the phenotypes of innate and adaptive immune cell types^19^. Because natural receptor domains achieve functional diversity through varied combinations of signaling motifs, we hypothesized that synthetic recombination of these motifs could polarize macrophages to diverse functional states. We further reasoned that a quantitative model that relates signaling motif combination to macrophage polarization would enable rational programming of macrophage phenotype.

We previously developed a synthetic cytokine receptor (SCR), which constitutively activates cell signaling cascades via short modular signaling motifs (∼16 amino acids, including a 4 to 8 amino acid consensus binding motif and flanking amino acids) in the receptor’s intracellular signaling domain. SCR signaling is cytokine-independent, activating signals in the absence of external input. SCRs with varied signaling motifs produce diverse chimeric antigen receptor (CAR) T cell states with unique survival, proliferation, differentiation, and cytotoxicity^20^.

Here, we extend the use of SCRs to program user-defined macrophage polarization states. By recombining signaling motifs in the SCR signaling domain, we generate a broad spectrum of synthetic macrophage polarization states, encompassing pro-inflammatory, anti-inflammatory, and highly phagocytic phenotypes. We find that expression of select SCRs can improve the targeted phagocytosis of cancer cells by CAR-macrophages *in vitro* and *in vivo*. We find that a two-state model describes the motif-to-phenotype relationship and use this model to rationally design an SCR whose expression drives macrophages to a state that is highly phagocytic, but also maintains a pro-inflammatory profile during a long-term CAR-macrophage killing assay. Together, these results establish a modular and programmable strategy for interrogating macrophage polarization and provide a foundation for the rational engineering of macrophage functions for therapeutic applications.

## Results

### Synthetic cytokine receptors drive ligand-independent macrophage polarization

Cytokine binding to natural receptors induces receptor complex assembly and activates signaling cascades. The specific signaling pathways activated are determined by the signaling motifs present on the receptors’ intracellular domain (ICD)^21^. To achieve cytokine-independent activation of user-defined signaling, we created synthetic cytokine receptors (SCRs) (Fig. 1a). A single lentiviral vector encodes both the SCR and green fluorescent protein (GFP) (Fig. 1a). Each SCR contains three domains: an extracellular coiled-coil that facilitates homodimerization, a transmembrane domain (TMD) positioned for maximal signaling^20,22^, and the gp130 box domain, which recruits Janus kinases (JAKs)^23^. These three domains are followed by signaling motifs in the distal region, which can be varied to alter cell signaling and thereby program cell phenotype. SCR0, which lacks signaling motifs, serves as a control.

**Figure 1.**
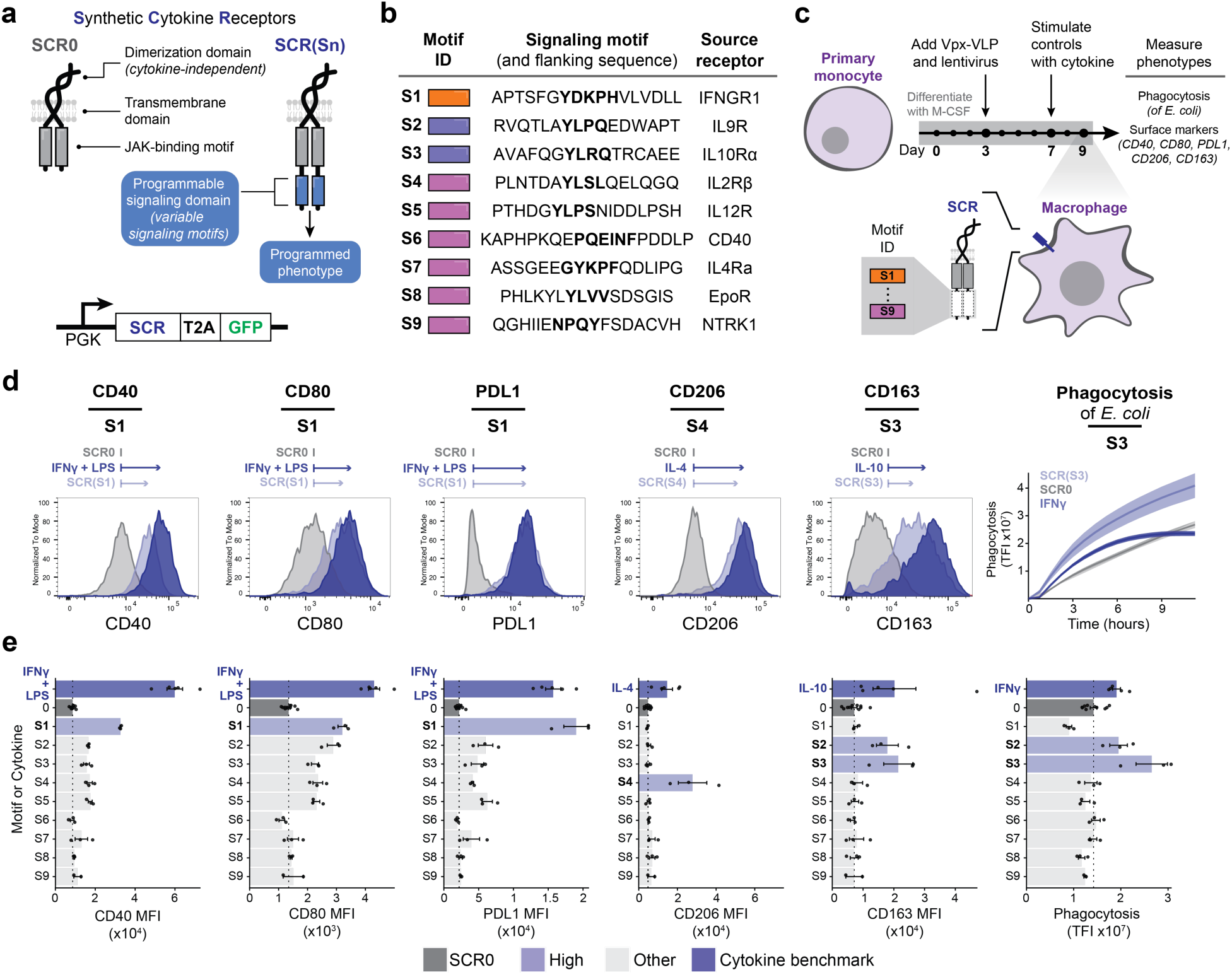
| Synthetic cytokine receptors drive ligand-independent macrophage polarization. **a,** Schematic depicting natural dimeric cytokine receptors (left) compared to the SCR platform (right). Also depicted is the SCR expression cassette driven by the phosphoglycerate kinase (PGK) promoter (bottom). The SCR and GFP are separated by a Thosea asigna virus 2A (T2A) self-cleaving peptide. SCR0, an SCR containing no motifs; SCR(Sn), an SCR containing a single synthetic signaling motif; TMD, transmembrane domain; JAK, Janus kinase. **b,** List of signaling motifs (S1 through S9), their full amino acid sequences, and the natural receptor they are derived from. The core sequence, known to bind to downstream effectors, is bolded. **c,** Schematic depicting the overall experimental design. Primary monocytes were differentiated and transduced with lentivirus containing an SCR. To benchmark against exogenous ligand treatment, control samples were treated (LPS, 10 ng/mL; IFN-γ, 50 ng/mL; IL-10, 50 ng/mL; IL-4, 50 ng/mL; GM-CSF, 50 ng/mL) 48 hours before phenotype measurement. Effects of each receptor were measured using surface marker flow cytometry and phagocytosis of pHrodo-Red-labelled *E.* c*oli*. **d,** Representative flow cytometry histograms and *E.* c*oli* phagocytosis of pHrodo-Red-labelled *E.* c*oli* measured by red total intensity (TFI) for macrophages expressing an SCR containing no motifs (SCR0; grey), macrophages expressing an SCR containing the indicated motif (light blue), and macrophages treated with the indicated stimulation (dark blue). For phagocytosis, data are mean ± s.e.m. **e,** Quantification of mean fluorescence intensity (MFI) or phagocytosis of pHrodo-Red-labelled *E.* c*oli* measured by red total intensity (TFI) for macrophages treated with the selected stimulation (dark blue; *n* = 5), macrophages expressing an SCR containing no motifs (grey; *n* = 10), or any single-motif SCR (light grey; *n* = 3). SCRs displaying an increase in the measured phenotype are indicated (light blue). Data are mean ± s.e.m. MFI, mean fluorescence intensity; TFI, total fluorescence intensity.

To construct SCRs with varied signaling, we selected nine signaling motifs from natural receptor ICDs (Fig. 1b). Each motif has binding or signaling activity that has previously been documented. Included are signaling motifs from the IL-10 receptor (IL-10R), which drives anti-inflammatory polarization, and the IFN-γ receptor (IFNGR1), which drives pro-inflammatory polarization^6^. We hypothesized that SCRs containing these motifs would drive distinct macrophage polarization states. To test this hypothesis, we constructed nine SCRs, each containing one of the motifs (S1 through S9) (Fig. 1b). We then co-transduced human primary monocyte-derived macrophages with lentivirus encoding the nine SCRs (Fig. 1c) and virus-like particles containing the protein Vpx, which enables efficient myeloid cell transduction^24–26^ (Extended Data Fig. 1b). To assess the effect of each SCR on macrophage polarization, we used flow cytometry to measure expression of select surface proteins that serve as macrophage polarization markers (CD40, CD80, PDL1, CD206, and CD163). *E. coli* labeled with a pH-sensitive dye (pHrodo) that fluoresces in the phagolysosome was used in live-cell imaging experiments to assess the effects of the SCRs on phagocytosis (Fig. 1c). We benchmarked the SCR-driven effects against 48-hour treatments with exogenous ligands (LPS and IFN-γ, IFN-γ, IL-10, IL-4, and GM-CSF) known to drive macrophage polarization into distinct states^6^ (Extended Data Fig. 1e).

For each of the measured surface markers, at least one motif increased expression to levels comparable to ligand stimulation (Fig. 1d and Fig. 1e). For example, SCR(S1), which contains the core amino acid “YDKPH” sequence from IFNGR1, increased CD40, CD80, and PDL1 expression (Fig. 1e). In contrast, SCRs containing motifs S2 or S3, each with a core “YxxQ” sequence, increased both CD163 expression and *E. coli* phagocytosis (Fig. 1e).

As expected, IL-4 treatment resulted in expression of CD206. However, motif S7, a reported STAT6-binding motif from the IL-4 receptor α chain^27^, did not increase CD206 expression. To investigate whether increasing the copy number of the S7 motif on the SCR would proportionally enhance CD206 expression, we expressed SCRs containing either one or three copies of the S7 motif. Three copies of S7 reached CD206 levels comparable to IL-4 stimulation (Extended Data Fig. 1c), suggesting that copy number can tune phenotype. Additionally, the S4 motif, a STAT5-binding motif from the IL-2 receptor β chain (IL-2Rβ)^28^, substantially increased CD206 expression, suggesting that STAT5 activation may also drive CD206 expression. Thus, the SCR can be used to drive specific phenotypes of interest, such as surface marker expression, through distinct signaling motifs.

Expression of SCR0 (the dimerization and JAK-binding scaffold alone) drove a modest increase in several measured phenotypes (Extended Data Fig. 1f). However, signaling motifs (Fig. 1e) and exogenous ligands (Extended Data Fig. 1f) drove significantly larger changes. Together, these results demonstrate that SCRs drive ligand-independent macrophage polarization into motif-dependent states distinguished by surface markers and phagocytosis.

### SCRs with signaling motifs from natural receptors mimic cytokine-induced macrophage polarization

The motifs S1 through S9 are small fragments of natural receptors, and our initial characterization of SCR-induced polarization investigated a narrow set of phenotypes. This raises two questions. First, can these signaling motifs recapitulate cytokine-induced macrophage polarization? Second, how do cytokine-induced and SCR-induced changes compare for more global phenotypic readouts like transcriptional state?

To address these questions, we compared the transcriptional programs induced by SCRs with those induced by cytokines that activate the native receptors from which the motifs were derived (Fig. 2a). We focused our inquiry on IFNγ and IL-10, which polarize macrophages into well-defined pro-inflammatory and anti-inflammatory states, respectively^4,29–31^. As in Fig. 1c, we transduced macrophages to express SCRs containing phosphotyrosine signaling motifs from either IFNGR1 (S1) or IL-10Rα (S3) (Extended Fig. 2b). We then performed bulk RNA sequencing (RNA-seq) on these macrophages and on macrophages treated with IFNγ or IL-10 for 48 hours. Cytokine-stimulated macrophages were also transduced with SCR0 to control for lentiviral transduction effects (Extended Fig. 2b). Monocytes from three different donors were used to assess donor variability (Extended Fig. 2a—b).

**Figure 2.**
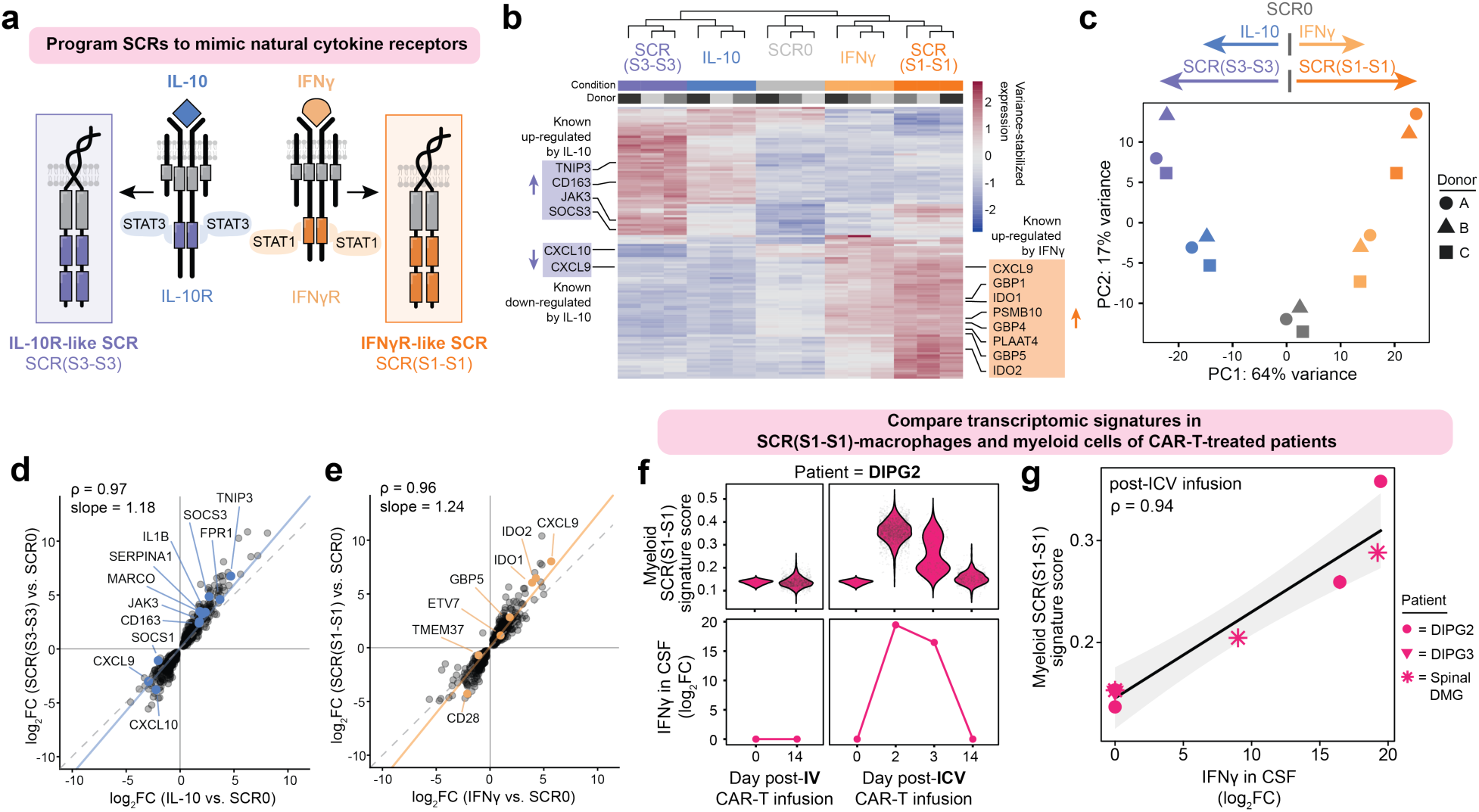
| Synthetic cytokine receptors mimic cytokine-driven macrophage polarization states. **a,** Schematic depicting the SCRs containing signaling motifs (S3 and S1) derived from the ICD of either the natural IL-10 receptor (IL-10R) or the natural IFN-γ receptor (IFN-γR), respectively. Motif sequences are indicated in Fig. 1b. **b,** Heatmap showing variance-stabilized expression of the top 50 differentially expressed genes across all sample comparisons (ranked by Benjamini–Hochberg–adjusted *p*-value). Columns and rows were hierarchically clustered using Euclidean distance. Genes known to be regulated in macrophages by either IL-10 or IFNγ are labeled. **c,** Principal component analysis (PCA) of RNAseq data from all donors and conditions. Donor is indicated by shape. **d,** Scatter plot of log_2_ fold changes for genes differentially expressed (padj < 0.05) in both IL-10 vs. SCR0 and SCR(S3-S3) vs. SCR0 comparisons (*n* = 773). Spearman correlation coefficient (ρ = 0.97) was calculated. The slope was estimated using linear regression constrained through the origin (green; slope = 1.18). Genes known to be regulated by IL-10 are indicated (blue). FC, fold change. **e,** Scatter plot of log_2_ fold changes (log_2_FCs) for genes differentially expressed (padj < 0.05) in both IFN-γ vs. SCR0 and SCR(S1-S1) vs. SCR0 comparisons (*n* = 667). Spearman correlation coefficient (ρ = 0.96) was calculated using the log_2_-fold-change (log_2_FC) values. The slope was estimated using linear regression constrained through the origin (green; slope = 1.24). Genes known to be regulated by IFN-γ are indicated (orange). FC, fold change. **f,** The SCR(S1-S1) transcriptional signature score of myeloid cells (above) and measured IFNγ levels (below) in the CSF of a patient with DIPG before and after either IV or ICV infusion of CAR-T therapy (data from Majzner and Ramakrishna et al, 2022^34^). FC, fold change; CSF, cerebrospinal fluid; DIPG, diffuse intrinsic pontine glioma; IV, intravenous; ICV, intracerebroventricular. **g,** Correlation between CSF levels of IFNγ and the SCR(S1-S1) transcriptional signature of myeloid cells from all patients in Majzner and Ramakrishna et al, 2022,^34^ for which these measurements were collected on the same day. Spearman correlation coefficient (ρ = 0.94) was calculated. FC, fold change; CSF, cerebrospinal fluid; DIPG, diffuse intrinsic pontine glioma; DMG, diffuse midline glioma.

SCR(S1-S1), SCR(S3-S3), IL-10, and IFNγ each changed gene expression relative to SCR0 (Fig. 2b). Hierarchical clustering showed that samples clustered by stimulation condition rather than donor (Fig. 2b). Consistent with their motif origins, macrophages expressing SCR(S3-S3) clustered with IL-10-treated macrophages, and macrophages expressing SCR(S1-S1) clustered with IFNγ-treated macrophages (Fig. 2b). The top differentially expressed genes (DEGs) included many DEGs shared between conditions. IL-10-stimulated and SCR(S3-S3) macrophages shared canonical IL-10-responsive genes (e.g., TNIP, CD163, JAK3, SOCS3) (Fig. 2b)^32^. Similarly, several IFNγ-responsive genes (e.g., CXCL9, GBP1, IDO1, GBP4, GBP5, IDO2) were shared between the IFNγ-treated and SCR(S1-S1) macrophages^31,33^ (Fig. 2b). These known cytokine-regulated genes also appeared in the top 10 DEGs across all samples, indicating strong activation of general macrophage polarization pathways (Extended Data Fig. 3a).

Principal component analysis showed a clear separation of cytokine-treated macrophages from the SCR0 control (Fig. 2c). Macrophages engineered with SCR(S1-S1) and SCR(S3-S3) projected in the same direction of PC1 as their cognate cytokines, but with greater displacement from the control (Fig. 2c). This observation led us to hypothesize that signaling motifs were driving similar transcriptional programs as their cognate cytokines, but to a stronger degree. Indeed, we observed a strong positive correlation (ρ = 0.96) in DEGs shared between the SCR(S3-S3) versus SCR0 comparison, and the IL-10 versus SCR0 comparison (Fig. 2d). The slope of the linear fit was greater than 1 (1.18), indicating a higher magnitude of differential expression in SCR(S3-S3)-expressing macrophages compared to IL-10-treated macrophages. We found a similarly positive correlation (ρ = 0.97) and slope (1.24) for DEGs shared between the SCR(S1-S1) versus SCR0 comparison and the IFNγ versus SCR0 comparison (Fig. 2e). In both cases, genes known to be regulated by IL-10 or IFNγ were predominantly positioned above the line of concordance, indicating stronger differential expression in SCR(S3-S3) and SCR(S1-S1) compared to cytokine-stimulated controls (Fig. 2d—e). Collectively, these results demonstrate that SCRs can mimic and exceed the transcriptional effects of cytokines at typically used concentrations (50 ng/mL).

### Motif S1 induces a myeloid activation state associated with CAR T cell activity in patients

The RNAseq data demonstrate that SCRs with signaling motifs from cytokine receptors can induce transcriptional features similar to those induced by cytokine treatment *in vitro*. We next investigated whether the SCR-driven transcriptional signatures reflect transcriptional signatures observed in clinical settings. A previous study included single-cell RNA sequencing (scRNA-seq) of cerebrospinal fluid (CSF) samples collected from four patients undergoing CAR T cell therapy for diffuse midline glioma^34^. This study identified an interferon response signature in myeloid cells that emerged in the days immediately following intracerebroventricular (ICV) CAR T cell infusion in a patient (DIPG2) who experienced improved neurological symptoms and a 27% reduction in tumor volume^34,35^. Patient DIPG2 also showed increases in CSF cytokines, including IFNγ and CXCL9, following ICV CAR T cell infusion. To determine if SCR(S1-S1) drives macrophages to a state reflective of this clinical myeloid response to CAR T cell therapy, we calculated an SCR(S1-S1) signature score for myeloid cells in the scRNA-seq dataset. The signature score quantifies the transcriptional similarity between patient myeloid cells and SCR(S1-S1) macrophages. In patient DIPG2, the SCR(S1-S1) signature was mostly highly enriched immediately following ICV CAR T cell infusion and subsequently decreased over the following days (Fig 2f, upper panel). This signature score closely paralleled the concentration of IFNγ measured in the patient’s CSF (Fig 2f, lower panel). To generalize this observation, we analyzed all time points across patients for which IFNγ measurements and CSF cells were collected on the same day. We observed a strong correlation (ρ = 0.94) between CSF IFNγ measurements and the myeloid SCR(S1-S1) gene signature score (Fig. 2g). These results demonstrate that the SCR(S1-S1)-driven transcriptional program observed *in vitro* is similar to the IFNγ-driven myeloid transcriptional programs observed in CSF following ICV CAR T infusion in a treatment-responsive patient. The alignment of SCR-macrophage transcriptomics with patient-identified myeloid signatures suggests programmed cells may serve as a model to understand the mechanistic implications of these cell types in patients.

### Signaling motif combinations enable navigation of diverse macrophage polarization states

We next asked if combining multiple motifs in one SCR can achieve unique polarization states not achieved using single cytokine stimulations. In contrast to the classical “M1/M2” paradigm, transcriptomic analyses have shown that polarized macrophages exist along a polarization spectrum, shaped by diverse stimuli^5^. Thus, we sought to explore whether expression of SCRs with several motifs would access a spectrum of polarization states (Fig. 3a). We generated a library of 131 unique SCRs (Fig. 3b) containing zero (n=1), one (n=9), two (n=81), or three (n=40) of the nine signaling motifs (Fig. 1b). With these 131 SCRs, we performed an arrayed experiment to assess each SCR’s effect on macrophage polarization (Fig. 3c). To contextualize these effects, we compared SCR-induced phenotypes to untransduced macrophages, SCR0-expressing macrophages, and macrophages stimulated with exogenous factors (IL-10, IFNγ, LPS + IFNγ, IL4, or GM-CSF).

**Figure 3.**
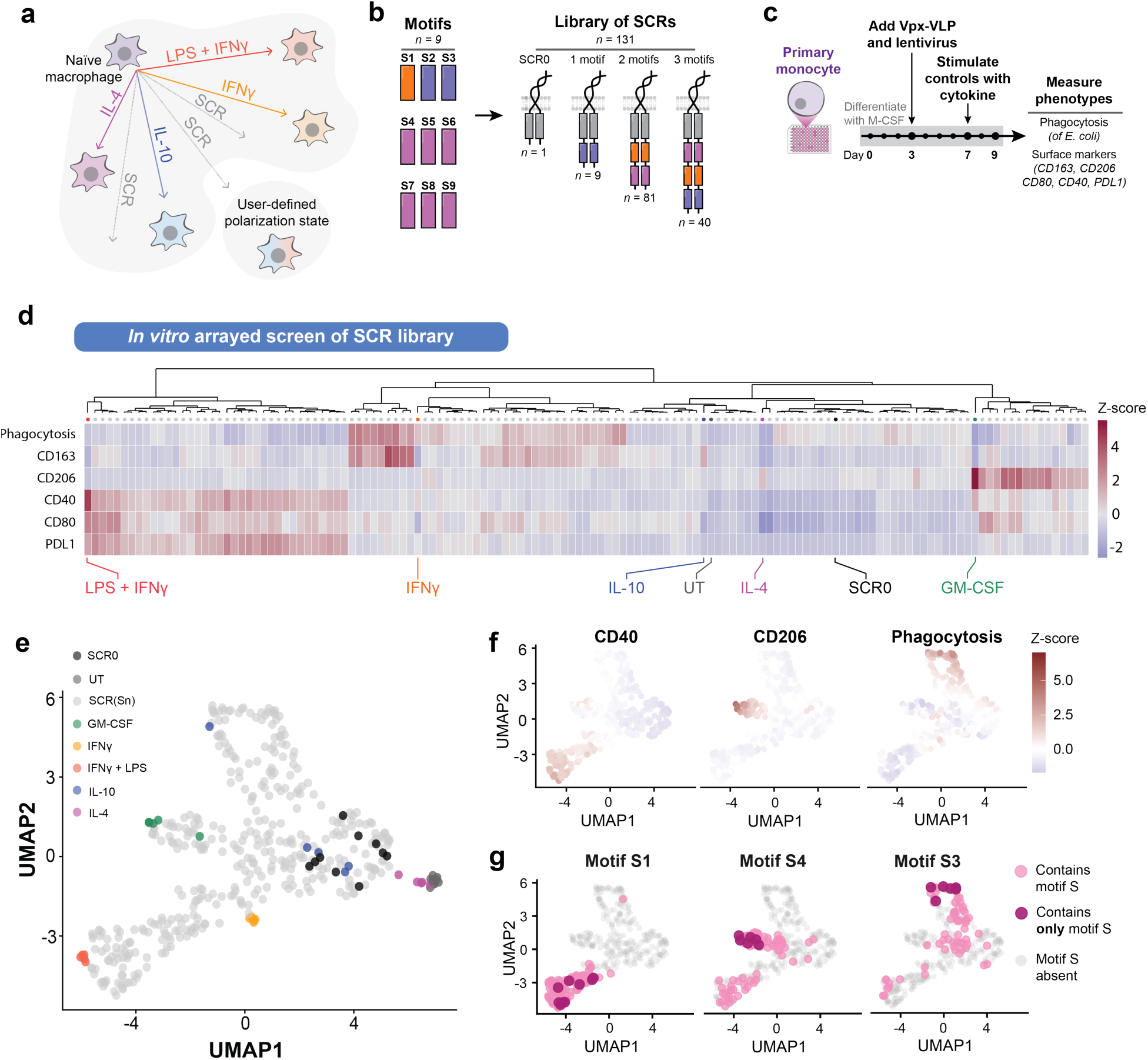
| Combining motifs on the SCR reveals a navigable landscape of primary macrophage polarization states. **a,** Schematic depicting the utility of recombining signaling motifs with SCRs to drive macrophages toward cytokine-induced polarization states, intermediate states, or user-defined states. **b,** Schematic of the combinatorial library of SCRs containing zero (SCR0; *n* = 1), one (*n* = 9), two (*n* = 81), or three (*n* = 40) signaling motifs (S1 through S9) from the motif library shown in Fig. 1b. **c,** Schematic of overall experimental design. **d,** Heatmap of macrophage phenotypes for all tested SCR constructs, cytokine stimulation controls, and transduction controls. Samples were hierarchically clustered using Euclidean distance based on z-scores of measured phenotypes. Dot color indicates sample type: macrophages stimulated with cytokines (LPS + IFN-γ, red, n = 5; IFN-γ, orange, n = 5; IL-10, blue, n = 5; IL-4, purple, n = 5), macrophages expressing SCR0 (black, n = 10), or macrophages expressing SCR constructs from the combinatorial library (light grey, n = 3 per SCR). *E. coli* phagocytosis was quantified using red integrated intensity measured 5.5 h after assay initiation. Sn, signaling motif. **e,** UMAP projection of all samples, including replicates. Controls are colored as in **d**. **f,** UMAP projection showing only samples expressing SCR constructs from the combinatorial library, colored by z-score for CD40, CD206, or *E. coli* phagocytosis. **g,** UMAP projection colored by motif presence for selected motifs (S1, S3, and S4). Pink indicates SCRs containing the indicated motif, while maroon indicates SCRs containing only that motif in one, two, or three copies. S, signaling motif.

The SCR library induced a broad spectrum of macrophage polarization states with distinct phagocytosis and expression of CD163, CD206, CD40, CD80, and PDL1 (Fig. 3d). This spectrum included both states that mimicked single cytokine treatments and states not achieved with cytokine stimulation. We used hierarchical clustering (Fig. 3d) and dimensionality reduction (Fig. 3e—g) to summarize SCR-induced polarization. UMAP embedding of all normalized surface marker and phagocytosis measurements showed that exogenous ligand treatments occupy distinct regions of the map, while SCR-driven phenotypes occupy the intervening phenotypic space (Fig. 3e). Five distinct functional clusters emerged. First, a CD40^high^ PDL1^high^ region occupied by LPS + IFNγ-stimulated macrophages (Fig. 3f). Second, a CD206^high^ area populated by GM-CSF-stimulated macrophages (Fig. 3f). Third, a region containing SCR-expressing samples with elevated phagocytic activity (Phago^high^) (Fig. 3f). Fourth, a baseline cluster containing SCR0 controls and SCRs with motif combinations that produced no significant phenotypic shifts (Fig. 3e). Lastly, a cluster of untransduced or IL-4-treated macrophages (Fig. 3e). The measured phenotypes were variably correlated with each other, with CD40 and PDL1 showing the strongest correlation (r = 0.86) (Extended Data Fig. 4b). Unexpectedly, CD80, a canonical pro-inflammatory marker, was expressed in multiple clusters, demonstrating that CD80 expression can occur outside strictly pro-inflammatory contexts (Extended Data Fig. 4c).

We analyzed how motifs were distributed across UMAP space to determine each motif’s contribution to macrophage polarization. SCRs containing solely the S1 motif (derived from IFNGR1) populated the CD40^high^PDL1^high^ region (Fig. 3g). This S1-driven effect dominated regardless of the presence or number of non-S1 motifs within the SCR (Fig. 3g). In contrast, motif S4 (derived from IL2Rβ) drove macrophages to the CD206^high^ region, while S2 (derived from IL9R) and S3 (derived from IL10Rα) drove macrophages to the Phago^high^ region. Unlike S1, the effects of S4, S2, and S3 were not dominant and could be modulated by the presence of other motifs within the SCR (Fig. 3g, Extended Data Fig. 4d). Motif position within the ICD did not affect phenotype. SCRs with reversed motif orders (e.g. ‘S1-S2’ vs. ‘S2-S1’) produced similar phenotypes (Extended Data Fig. 4e). Additionally, motif copy number impacted phenotype differently for each motif and phenotype (Extended Data Fig. 4f). For example, one copy of S1 drove maximal PDL1, CD40 and CD80 expression, while S2 increased phagocytosis proportionally with copy number (Extended Data Fig. 4f). Collectively, these findings are consistent with the macrophage polarization spectrum model and demonstrate that the complex phenotypic landscape can be navigated by varying SCR motif combinations.

### SCRs containing motifs S2 or S3 improve the anti-tumor activity of CAR macrophages

We next sought to evaluate whether SCRs could improve the potential therapeutic functions of macrophages. Chimeric antigen receptor-expressing macrophages (CAR-Ms) have emerged as promising immunotherapeutics for a range of diseases, including cancer, cardiac fibrosis, and bacterial infections^36–39^. The efficacy of CAR-Ms in these therapies will depend on factors like inflammatory state and phagocytic capacity.

Macrophages expressing SCRs with S2 or S3 motifs showed increased phagocytosis of *E. coli* (Fig. 1d and Fig. 3d). We therefore asked whether SCRs containing these motifs would augment CAR-M-mediated phagocytosis of cancer cells. To answer this, we co-expressed each of four SCRs that in increased *E. coli* phagocytosis— SCR(S2-S2-S2), SCR(S3-S3), SCR(S5-S3), SCR(S2-S3)—with a second-generation αHER2-41BBζ chimeric antigen receptor (CAR) in primary human macrophages (Fig. 4a). We then co-cultured these CAR-SCR-Ms with BT474 breast cancer cells, which express high levels of the antigen human epidermal growth factor 2 (HER2) (Fig. 4b). Macrophages were challenged with pHrodo-labeled BT474 cells three times over nine days at increasing effector-to-target ratios (Fig. 4b).

**Figure 4.**
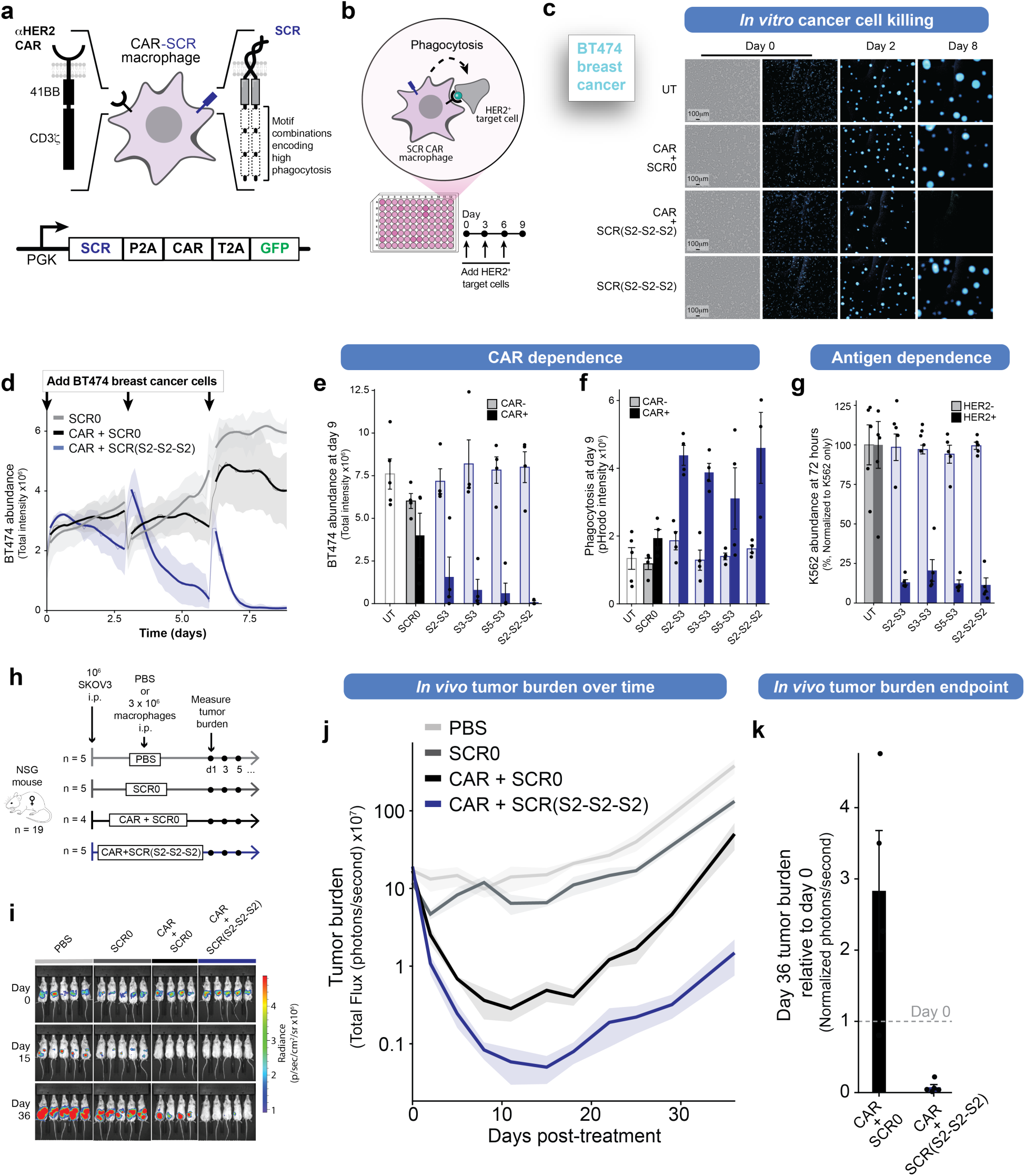
| SCR expression augments phagocytosis of cancer cells by CAR-Macrophages. **a,** Schematic depicting co-expression of a human epidermal growth factor (HER)2-specific chimeric antigen receptor (CAR) and an SCR in human primary macrophages. Also depicted is the co-expression cassette driven by the phosphoglycerate kinase (PGK) promoter (bottom). HER2, human epidermal growth factor; GFP, green fluorescent protein. **b,** Schematic depicting the experimental design for repeated challenge of αHER2-CAR macrophages with HER2⁺ cancer cells. HER2, human epidermal growth factor 2. **c,** Representative images of BT474 cell abundance over 9-days of repeated co-culture with macrophages. Macrophages were transduced with the indicated constructs. UT, untransduced. **d,** BT474 cell abundance over nine days of repeated co-culture with macrophages transduced with SCR0 (*n* = 4), CAR + SCR0 (*n* = 4), or CAR + SCR (S2-S2-S2) (*n* = 3). BT474 cell abundance was measured as total BFP fluorescence intensity using live-cell imaging. Faint lines represent the raw mean value. Darker lines represent the smoothed curve for each challenge period. Data are mean ± s.e.m. **e,** BT474 cell abundance after nine days of repeated co-culture with macrophages transduced with SCRs containing motifs that drove high phagocytosis in Fig. 3d, with (light) or without (dark) co-expression of a CAR. Data are mean ± s.e.m. **f,** Quantification of pHrodo signal intensity after nine days of repeated co-culture with macrophages transduced with SCRs containing motifs that drove high phagocytosis in Fig. 3d, with (light) or without (dark) co-expression of a CAR. Data are mean ± s.e.m. **g,** Quantification of relative K562 growth at the end of a 3-day co-culture between CAR-SCR-Ms and wild-type K562 cells (light) or HER2^+^ K562 cells (dark). K562 cell growth is the normalized growth of K562s in the absence of macrophages (K562 only). Data are mean ± s.e.m. **h,** Schematic depicting the *in vivo* experimental design. 1x10^6^ SKOV3 cells expressing firefly luciferase (ffluc) were injected i.p. into each mouse. Mice were then treated i.p. with PBS (grey, *n* = 5) or 3x10^6^ macrophages expressing SCR0 (grey, *n* = 5), αHER2-CAR + SCR0 (light blue, *n* = 4), or αHER2-CAR + SCR(S2-S2-S2) (dark blue, *n* = 5). Tumor burden was measured using bioluminescent imaging (BLI). i.p., intraperitoneal. **i,** Representative images from the experiment depicted in **f** at day 0, day 15, and day 36 after tumor injection. BLI, bioluminescent imaging. **j**, Tumor burden measured as total flux (photons s⁻¹) of luciferase signal. **k,** Tumor burden at day 36 post treatment, normalized by the day 0 luciferase signal.

CAR-Ms expressing any of the four SCRs significantly reduced BT474 survival compared to CAR-SCR0-Ms (Fig. 4c–e). Correspondingly, pHrodo signal increased in these samples, demonstrating that phagocytosis contributed to this enhanced killing (Fig. 4f). Importantly, the decrease in BT474 survival was dependent on expression of the CAR, as SCR expression alone did not cause a reduction in BT474 survival (Fig. 4c, e). To additionally test whether CAR-SCR-M-mediated killing depends on the presence of target antigen, we co-cultured CAR-SCR-Ms with either wild-type (WT) K562 cells or K562 cells overexpressing HER2 (HER2^+^) for 72 hours. CAR-SCR-Ms killed HER2^+^ K562s but not WT K562s (Fig. 4g). Together, these data demonstrate that SCRs can augment antigen-dependent CAR-mediated killing of cancer cells *in vitro* without increasing off-target killing.

We further evaluated the anti-tumor activity of CAR-SCR-Ms *in vivo* using a SKOV3 human ovarian cancer xenograft model. SKOV3s, which naturally express HER2, were injected intraperitoneally (i.p) into NOD.Cg-Prkdc^scid^ Il2^rgtm1Wjl/SzJ^ (NSG) mice. Mice then received a single i.p. injection of either saline (PBS), SCR0-Ms, CAR-SCR0-Ms, or CAR-SCR(S2-S2-S2)-Ms (Fig. 4h). Both CAR-M treatments significantly reduced tumor burden compared to PBS or SCR0-M controls (Fig. 4i—j). Strikingly, CAR-SCR(S2-S2-S2)-Ms reduced tumor burden 30-fold compared to CAR-SCR0-Ms at day 36 post-injection (Fig. 4k). SCRs thus improve CAR-M therapeutic function in a disease-relevant context. Because SCRs enhance CAR-mediated phagocytosis without altering the CAR itself, SCRs can be readily combined with existing CARs to create more effective CAR-M therapies.

### Signaling motifs predictably alter macrophage phenotypes to enable programmable control of macrophage polarization

While SCRs with unique combinations of one to three signaling motifs generated a wide range of macrophage polarization states, these SCRs represent only ∼1.78% of the 7,372 possible SCRs containing up to four motifs. Future applications may require expanding the motif selection and increasing the number of motifs per SCR. As this combinatorial space grows, experimentally testing all SCRs becomes impractical, necessitating computational models to predict optimal SCR designs from limited experimental data^40^. Neural networks can predict the effects of signaling motifs on CAR T cell phenotype^20,41^, but these models require more data than is present in our macrophage array dataset. Thus, we sought a data-efficient model that connects the choice of signaling motifs to measurable phenotypes such as cellular activity (e.g., phagocytosis) or expression of a protein.

Two-state models have relatively few parameters and have been used to describe various biological phenomena, such as protein folding reactions or receptor-ligand interactions^42,43^. We employed a two-state model to predict how SCR signaling motifs program macrophage polarization (Fig. 5a, Extended Data Fig. 7). Because motif position does not alter measured phenotypes (Extended Data Fig. 4e), the model assumes motif effects are position-independent. This model also assumes that for each phenotype, cells can exist in an ON state or an OFF state whose relative populations are defined by the polarization constant *K* (Fig 5a). Without SCR signaling, the relative populations of the ON and OFF states are defined by the intrinsic polarization constant, *Ki* (Fig. 5b). Each motif added to the SCR shifts these populations of the ON and OFF states for a given phenotype. The effect of each motif on each phenotype is described by motif-dependent polarization constant*s, Ks* (*s*, signaling motif) (Fig. 5b).

**Figure 5.**
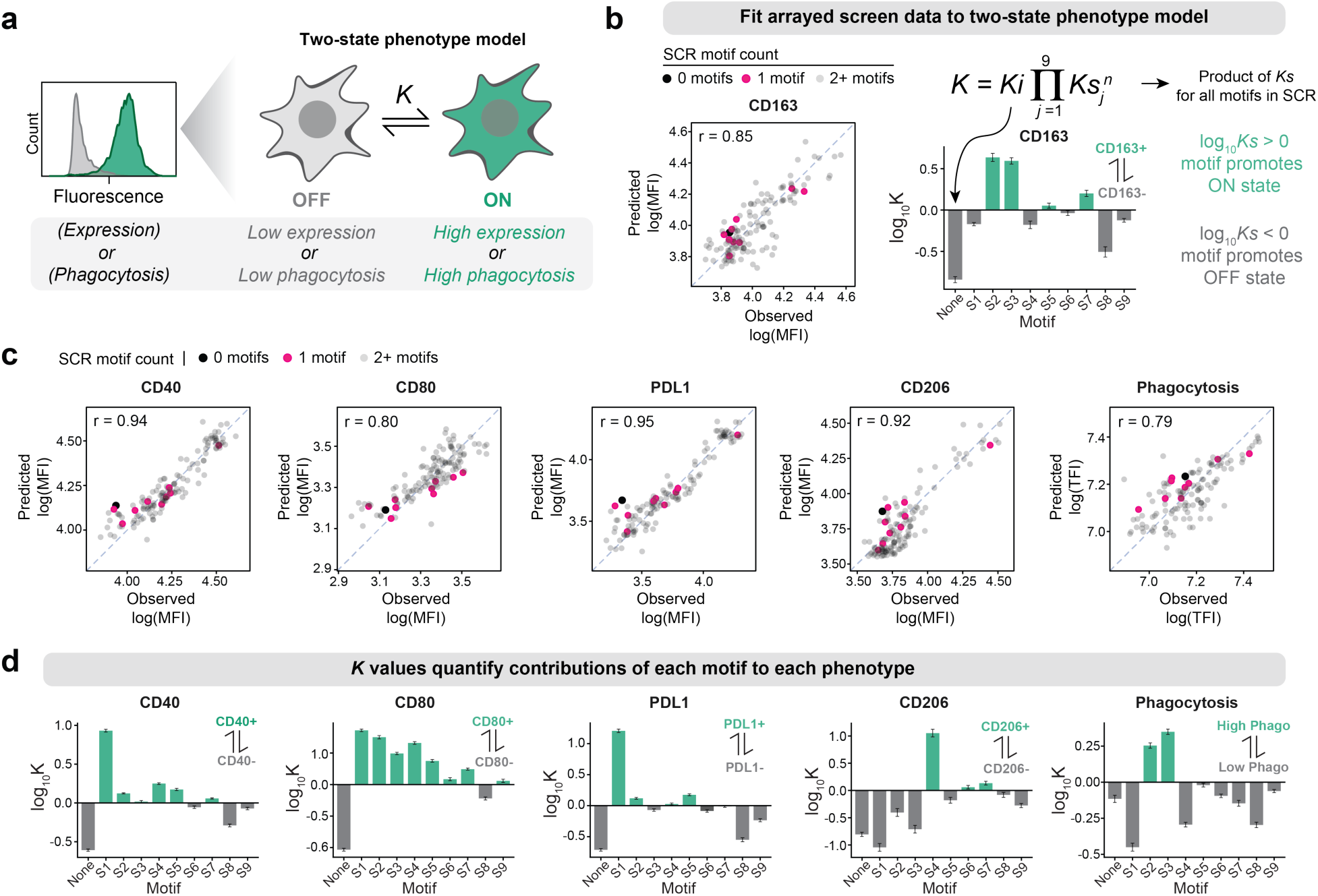
| A two-state model describes the contribution of each signaling motif to each measured phenotype. **a,** Schematic of the two-state phenotype model that describes polarization of macrophages to ON and OFF phenotypic states. *K* represents the relative population of cells in the ON and OFF states. **b,** Observed and predicted CD163 MFI and *K* values estimated from the model. MFI, mean fluorescence intensity. **c,** Observed vs. Predicted scatterplots for each measured phenotype (MFI of surface marker or phagocytosis of *E. coli*). Points are labeled by the number of motifs in the sample (no motifs, black; single motif, pink; more than 1 motif, grey). Observed data are averaged per motif combination. MFI, mean fluorescence intensity; TFI, total fluorescence intensity. **d,** Calculated *Ks* parameter values for each motif for each phenotype. Positive *Ks* values (green) indicate a shift toward the positive state for the indicated phenotype, while negative *Ks* values (grey) indicate a shift toward the negative state for the indicated phenotype. Data are fitted parameters ± s.d. of cross-validation folds (*n* = 10).

We fit this two-state model to the array data (Fig. 3d) to estimate the effects of each motif (*Ks*) on each phenotype (Fig. 5b). The data were well-described by the model. We observed high correlation between predicted and observed values across phenotypes, including surface marker expression (CD163 (r = 0.85), PDL1 (r = 0.95), CD40 (r = 0.94), CD80 (r = 0.80), CD206 (r = 0.92)), and phagocytic activity (r = 0.79) (Fig. 5b—c). Estimated *Ks* values reflect the impact of each motif on phenotype. These values were consistent with the prior observation that motif S1 drove macrophages to a CD40^high^ state (log_10_*Ks* = 1.01) (Fig. 5d), motifs S2 and S3 drove macrophages to the Phago^high^ state (log_10_*Ks*(S2) = 0.32; log_10_*Ks*(S3) = 0.70), and motif S4 drove macrophages to a CD206^high^ state (log_10_*Ks*(S3) = 1.17) (Fig. 5d). These modeling results suggest that signaling motifs, ubiquitous elements in cytokine receptors that control macrophage phenotype, shift phenotype in a predictable manner.

### Modeling guides programming of signaling domains that polarize macrophages to a highly phagocytic and pro-inflammatory cell state

We sought to leverage the two-state model to engineer a macrophage with a unique polarization profile advantageous in a disease setting. One desirable macrophage state for anti-cancer therapy is one that both phagocytoses cancer cells efficiently and remodels the tumor microenvironment through pro-inflammatory actions^44^. However, within the experimentally tested SCRs, we noticed a tradeoff between the level of phagocytosis and the expression of pro-inflammatory markers (CD40 and CD80) by SCR-expressing macrophages (Fig. 6a). No SCR drove macrophages to a high CD40 and CD80 expression state while also maintaining enhanced phagocytosis (Fig. 6a). We hypothesized that the two-state model could be used to identify motif combinations that overcome this tradeoff—specifically, combinations that would drive macrophages to the CD80^+^/CD40^+^/phagocytosis^high^ quadrant (Fig. 6a).

**Figure 6.**
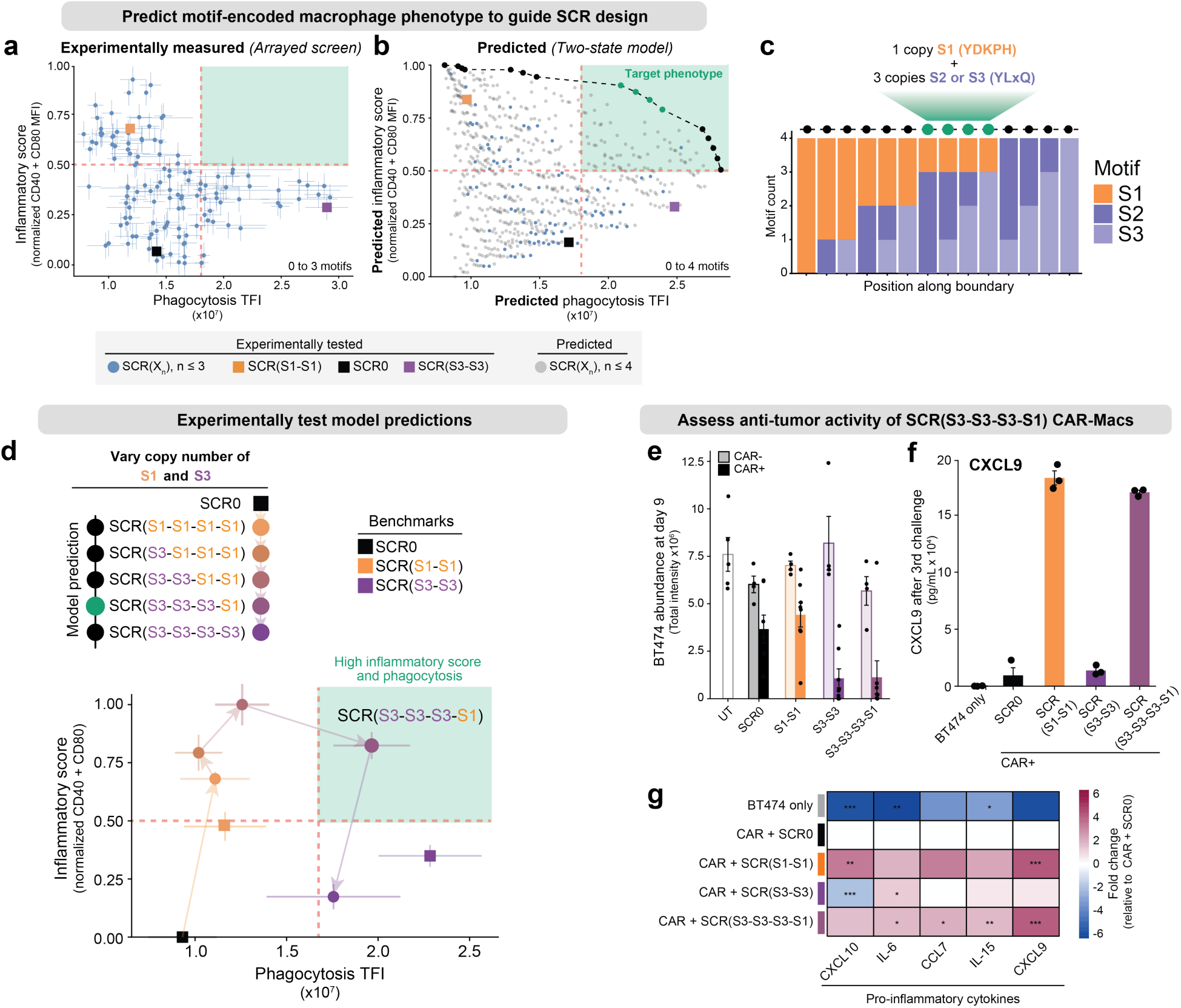
| Modeling guides rational SCR design to drive a simultaneously pro-inflammatory and phagocytic macrophage state. **a,** Scatterplot of phagocytosis of *E. coli* (Phagocytosis TFI) by inflammatory score (normalized sum of CD40 and CD80 MFI) for experimentally tested SCR-expressing macrophages. The dashed line indicates quadrants that can be drawn on top of the point distribution. MFI, mean fluorescence intensity; TFI, total fluorescence intensity. Data are mean ± s.e.m. **b,** Scatterplot of predicted phagocytosis of *E. coli* (Phagocytosis TFI) by inflammatory score (normalized sum of CD40 and CD80 MFI) for all SCRs containing between 0 and 4 motifs. SCRs that were previously tested experimentally are indicated (blue). SCRs predicted to drive macrophages to values along the boundary are indicated (black). SCRs predicted to drive macrophages to a simultaneously high inflammatory and high phagocytosis state are indicated (green). The dashed line indicates quadrants that can be drawn on top of the point distribution. MFI, mean fluorescence intensity; TFI, total fluorescence intensity. **c,** Total motif count for each motif along the boundary labeled in **b**. **d,** Scatterplot of measured phagocytosis by inflammatory score (normalized sum of CD40 and CD80 MFI) for macrophages expressing SCRs containing motif S1 and motif S3 at varying copy number ratios. Points are colored by the copy number of each motif present in the SCR (S1, orange; S3, purple). Data are mean ± s.e.m. **e,** BT474 cell abundance after nine days of repeated co-culture with macrophages transduced with controls or the model-guided motif combination (SCR(S3-S3-S3-S1)), with (light) or without (dark) co-expression of a CAR. Data are mean ± s.e.m. **f,** Concentration of CXCL9 in the supernatant at the end of the third challenge from the experiment in d. Data are mean ± s.e.m. **g,** Heatmap showing log_2_ fold-change in selected cytokine concentration in the supernatant at the end of the third addition of BT474 cells to CAR macrophages co-expressing the indicated SCR, relative to the CAR + SCR0 control. Only cytokines with minimal baseline secretion by BT474 cells alone are shown. Statistical significance was assessed using a two-sample t-test comparing replicate concentration values (n = 3) for each condition against the CAR + SCR0 baseline for each cytokine independently.

We used the fitted *Ks* values to predict CD40, CD80, and phagocytosis measurements for all possible SCRs containing up to four motifs. These predictions recapitulated the tradeoff between phagocytosis and CD40/CD80 expression for the SCR designs that were experimentally tested (Fig. 6b). However, we also found that several untested SCRs were predicted to drive macrophages to the CD40^+^/CD80^+^/Phago^high^ quadrant (Fig. 6b). SCRs along the trade-off boundary (black and green points) contained decreasing copies of motif S1 and increasing copies of motif S2 and S3, with the four SCRs predicted to simultaneously maximize CD40, CD80 and phagocytosis containing one S1 copy and three copies of either S2 or S3 (Fig. 6c).

To test these predictions, we constructed six, four-motif SCRs spanning all stoichiometric combinations of S1 and S3 (Fig. 6d). We benchmarked these against SCR(S1-S1) and SCR(S3-S3) from the earlier tested library (Fig. 3b). As expected, two of the three S1/S3 SCRs drove the S1-dominant phenotype (CD40^+^/CD80^+^/Phago^low^), while macrophages expressing one S1 copy and three S3 copies (SCR(S3-S3-S3-S1)) displayed simultaneously high phagocytosis and high CD40 and CD80 expression (Fig. 6d). To understand if the high phagocytosis and inflammatory phenotype generated by SCR(S3-S3-S3-S1) is maintained in the context of CAR-M, we followed the experimental design established in Fig. 4a and Fig. 4b. Expression of SCR(S3-S3-S3-S1) enhanced CAR-dependent killing of BT474 cells to a similar extent as SCR(S3-S3), with both constructs promoting enhanced killing compared to SCR0 (Fig. 6e).

We previously found that macrophages expressing SCR(S1-S1) mimic the polarization state induced by IFNγ at the transcriptional level (Fig. 2b) and that one of the top differentially expressed genes between SCR(S1-S1) and SCR0-expressing macrophages is CXCL9 (Fig. 2c, Extended Data Fig. 3a). The IFNγ/CXCL9 signaling program is known to promote anti-tumor immunity by encouraging cytotoxic T lymphocyte recruitment and effector differentiation^1^. It has also been shown that CXCL9 expression requires sustained IFNγ signaling in macrophages^45^, making the constitutive nature of SCR signaling potentially advantageous for driving CXCL9 expression and an anti-tumor response. To see if CXCL9 secretion is enhanced in CAR-Ms expressing SCR(S1-S1) over several days and if this effect is retained when expressing SCR(S3-S3-S3-S1), we collected supernatant at the end of the 9-day CAR-M co-culture experiment. We found a significant increase in the concentration of CXCL9 in the media when macrophages expressed SCR(S1-S1) compared to SCR0. This effect was not observed in macrophages expressing SCR(S3-S3) yet was retained in macrophages expressing SCR(S3-S3-S3-S1) (Fig. 6f). Macrophages expressing SCR(S3-S3-S3-S1) also exhibited greater secretion of CXCL10, CCL7, and IL-15 (Fig. 6g), other pro-inflammatory cytokines known to recruit a diversity of immune cells to the tumor microenvironment. Together, these data demonstrate that rational, model-guided SCR design can generate a macrophage with simultaneously enhanced phagocytosis and pro-inflammatory signaling. More broadly, this exemplifies the capacity of the SCR platform to fine-tune complex cellular phenotypes through model-guided receptor design.

## Discussion

Macrophages adopt a wide array of polarization states to carry out diverse functions. However, precise control over these states remains a challenge. Here, we leveraged synthetic signaling to systematically program human primary macrophage polarization. Using a two-state model to quantify the motif-to-phenotype relationship, we rationally designed a synthetic receptor that overcomes an observed trade-off between macrophage inflammatory markers and phagocytosis. We additionally showed that this fine-tuned phenotype persists in CAR-macrophages during a long-term co-culture with cancer cell targets. This highlights an important advantage of the motif recombination strategy: each motif has a unique effect on phenotypes like phagocytosis, CD80 expression, or CD40 expression. Recombination of motifs therefore allows multi-dimensional programming of phenotypes that are not inherently coupled.

Building on recent advances in engineered macrophage cell therapies, we used an SCR to enhance the killing of solid tumor cells by CAR-Ms. Further optimization of SCRs could increase CAR-M survival and persistence of these cells to increase anti-tumor efficacy. The wide range of SCR-driven phenotypes observed in our arrayed screen may also be useful in different therapeutic contexts. For example, phagocytic pro-inflammatory CAR-Ms may be useful in the treatment of immunosuppressive tumors^44^, while anti-inflammatory macrophages that release Secreted Phosphoprotein 1 (SPP1) may be useful in treating spinal injuries^47^.

The programmability enabled by signaling motif recombination offers a significant advantage over traditional use of exogenous cytokines. By combining signaling motifs from different receptors, we can generate unique signaling^20^—and corresponding macrophage phenotypes—that would be difficult or impossible to control using conventional cytokine stimulation. This programmability creates new possibilities for rational design of SCRs that induce desired macrophage phenotypic profiles for various therapeutic applications, as well as T cells and potentially other immune cells. Though this study focuses on the use of SCRs to polarize macrophages, we have also used SCRs to program T cell differentiation^20^. Considering the conservation of cell signaling machinery across cell types, we suspect that the utility of SCRs will extend to other immune cell types—such as natural killer cells and dendritic cells—and hematopoietic stem cells.

## Materials and Methods

### Study Design

This study was designed to determine if expression of synthetic cytokine receptors (SCRs) could drive polarization of macrophages. SCRs were cloned and expressed in human primary macrophages. Effects of SCR expression on several macrophage phenotypes was assessed using surface marker staining and flow cytometry or by co-culturing the macrophages with biological targets of phagocytosis followed by live cell imaging. Shifts in measured phenotypes were compared to those achieved when untransduced macrophages were stimulated for 48 hours with select exogenous ligands. To determine the transcriptional effects of expressing select SCRs in macrophages, we performed bulk RNA sequencing. To discover if SCRs could generate a wide range of macrophage phenotypes, we recombined 9 signaling motifs on the intracellular domain of the SCR to generate >100 SCRs and tested these using previously described methods. Data from this screen were used to fit a two-state equilibrium model and predict SCR designs to drive both a high inflammatory profile and high phagocytosis. To evaluate if SCRs associated with high phagocytosis could augment the targeted phagocytosis of cancer cells by CAR-Macrophages, we co-cultured macrophages expressing both an SCR and a CAR (SCR-CAR-Ms) with cancer cells expressing the CAR-targeting antigen and a fluorescent marker. Cancer cell number was monitored via live cell imaging. SCR-CAR-Ms were then tested for their ability to phagocytose cancer cells in mice using an ovarian tumor model. Numbers of biological and experimental replicates and statistical analyses are reported in the figure legends.

### Lentiviral vector construction

All DNA constructs were visualized using SnapGene software (v.8.1.1; Dotmatics). DNA fragments encoding the variable library parts (S1 through S9), SCR, T2A-GFP, and P2A-CAR were codon optimized for expression in human cells using ThermoFisher GeneArt’s website tool and synthesized by ThermoFisher GeneArt. All DNA fragments were isolated by agarose gel purification using a NuceloSpin Gel and PCR Clean-up Kit (Macherey-Nagel #740609.50). All cloning reactions were conducted using In-Fusion Snap Assembly (Takara Bio USA #638949). All plasmids were sequence verified using Sanger sequencing (MCLAB) before use.

Lentiviral vectors encoding the synthetic cytokine receptor were generated as described in Cho et al. (2025)^20^. The synthetic cytokine receptor (SCR) is a chimeric protein consisting of a signal peptide, myc tag, Put3 coiled-coil domain, erythropoietin receptor transmembrane domain, gp130 box motifs and disordered region (residue 642 to residue 725), and gp130 dileucine internalization motif (residue 780 to residue 796). This SCR sequence, along with a fragment encoding T2A-GFP were then cloned into the pHR-PGK (Addgene #79120) backbone. This pHR-PGK-SCR-T2A-GFP construct is indicated as the SCR containing no motifs (SCR0).

To integrate motifs into the SCR0 construct, motifs were restriction digested using flanking BfuAI sites, and fragments were cloned into the SCR0 construct. This cloning process resulted in the SCR backbone, followed by the motif sequence, in the same frame, separated by a (GS)_3_ linker sequence. Each motif contains the proper sequence to regenerate the BamHI restriction site for insertion of subsequent motifs.

For constructs containing both an SCR and a CAR, a T2A-αHER2-41BBζ CAR was inserted upstream of the P2A-GFP to create SCR-(MOTIF)_n_-T2A-αHER2-41BBζ-P2A-GFP. The P2A-CAR is a chimeric protein consisting of a P2A sequence, signal peptide, hemagglutinin (HA) tag, anti-HER2 4D5 single-chain variable fragment, CD8a hinge and transmembrane domain, 41BB-derived costimulatory domain, and CD3z signaling domain.

For *in vitro* and *in vivo* tracking of cancer cells, a dual BFP/firefly-luciferase lentiviral expression vector was constructed. Lenti-NLS-mTagBFP2 plasmid (Addgene #216128) was digested with PacI. A P2A and firefly luciferase (ffluc) sequence (from Addgene #115684) were PCR amplified and cloned into the linearized lenti-NLS-mTagBFP2 to create lenti-NLS-mTagBFP2-P2A-ffluc.

### Human primary cells

Human peripheral blood from healthy anonymous donors was purchased as apheresis leukopaks (STEMCELL Technologies #70500.1). Human primary monocytes were negatively selected from leukopaks using EasySep Human Monocyte Isolation Kit (STEMCELL Technologies #19359) according to the manufacturer’s instructions. Isolated monocytes were suspended in RPMI-1640 containing 20% DMSO and 20% FBS and cryopreserved in liquid nitrogen until use. Upon thawing, monocytes were centrifuged at 400xg for 4 minutes, and the supernatant was aspirated to remove DMSO. Monocytes were then cultured in Immunocult-SF Macrophage Medium (STEMCELL Technologies #10961) containing human recombinant M-CSF (STEMCELL Technologies #78057) at 50 ng/mL. Isolated monocytes were profiled by flow cytometry using αCD45, αCD14, and αCD16 antibodies as described in **Flow cytometry > Monocyte isolation profiling**. All tissue culture was performed under sterile conditions. All cells were kept in temperature-controlled incubators at 37°C with 5% CO_2_.

### Lentivirus production

Lentiviral supernatant was produced in the Lenti-X 293T packaging cell line (Clontech #11131D). 293T cells were grown to ∼70% confluency before transfection. VSVG-pseudotyped lentivirus was generated by transient transfection of 293T cells with lentiviral packaging plasmids pCMVdR8.91 and pMD2.G (Addgene #12259), using FuGENE HD (Promega #E2312) at a 3:1 DNA:FuGene ratio. Transfection solutions were diluted in Opti-MEM (Gibco, Opti-MEM) and incubated for 10 minutes before being added to the 293T cell culture.

#### In vitro experiments

For *in vitro* experiments, 293T cells were plated on TC-treated flat-bottom plates. 200ngs of total DNA was used for transfection in a 96-well format or 3.4μgs of total DNA in a 6-well format. Lentiviral supernatant was harvested at 48 hours, centrifuged to deplete cell debris, and used fresh. Transfections were conducted side-by-side to reduce sample-to-sample variability.

#### In vivo experiments

For *in vivo* experiments, 293T cells were plated on 10cm TC-treated dishes. 12μgs of total DNA was used for the transfection of each plate. The media was completely changed the following day. Lentiviral supernatant was harvested at 48 hours after media change, centrifuged to deplete cell debris, transferred to a new container, and concentrated using Lenti-X Concentrator (Takara #631232) using a 3:1 supernatant:LentiX ratio by volume. Concentrated lentivirus was stored in - 80°C for later use. Concentrated lentivirus stock was titrated using serial dilution and flow cytometry to optimize for transduction efficiency and transduced macrophage yield.

### Virus-like particle production

Virus-like particles containing Vpx protein were generated by transient transfection of 293T cells with pSIV-D3psi (Addgene #132928) and pMD2.G (Addgene #12259) using FuGENE HD (Promega #E2312) at a 3:1 DNA:FuGene (mass:volume) ratio. After 48 hours, lentiviral supernatant was collected, centrifuged to deplete cell debris, transferred to a new container, and concentrated using Lenti-X Concentrator (Takara #631232) using a 3:1 supernatant:LentiX ratio by volume and frozen at -80°C until use. Vpx^+^ VLPs were titrated using serial dilution and imaging of GFP positivity of human primary macrophages upon co-transduction with a GFP^+^ lentiviral vector to establish the lowest amount of VLPs that could be used for efficient transduction. Large volumes of stock Vpx^+^ VLPs were generated and aliquoted to reduce variability across experiments.

### Primary macrophage transduction

#### In vitro experiments

Macrophages for either flow cytometry (60E5 cells per well), phagocytosis of *E. coli* (25E4 cells per well), or bulk RNA sequencing (1E6 cells per well) were seeded onto a TC-treated 96-well plate, flat-bottom 96-well plate (Thermo Scientific, 167008), or TC-treated 6-well plate, respectively. Cells were cultured in serum-free macrophage media (STEMCELL Technologies, #10961) and differentiated with 50 ng/mL of M-CSF (STEMCELL Technologies, #78057) for 4 days before transduction. Cells were then co-transduced with both Vpx^+^ virus-like particle stock (described in **Virus-like particle production**) and lentiviral supernatant containing lentivirus with the construct of interest (described in **Lentivirus production**). Cells were incubated overnight. Lentiviral media was replaced the following morning with fresh, serum-free macrophage media (STEMCELL Technologies, #10961) containing M-CSF 50 ng/mL (STEMCELL Technologies, #78057). Cells were incubated at 37°C until endpoints were measured.

#### In vivo experiments

Primary monocytes were thawed and seeded (described in **Human primary cells**) onto 6-well Nunc dishes with an UpCell surface (Thermo Scientific). Four days after thaw, cells were co-transduced with both Vpx^+^ virus-like particle stock (described in **Virus-like particle production**) and concentrated lentivirus containing the construct of interest (described in **Lentivirus production**). Cells were incubated overnight. Lentiviral media was replaced the following morning with fresh, serum-free macrophage media (STEMCELL Technologies, #10961) containing M-CSF 50 ng/mL (STEMCELL Technologies, #78057). Cells were incubated for 6 days, with M-CSF being replenished after 3 days.

### Cytokine-mediated macrophage polarization

Cytokine-mediated polarization was performed as controls. The following cytokines or other factors were added (at the indicated final concentration) to the media, 48 hours before measurements were performed: LPS (10 ng/mL) (Sigma-Aldrich, L4391), IFN-γ (50 ng/mL) (STEMCELL Technologies, #78020), IL-4 (50 ng/mL) (Gibco, 200-04), IL-10 (50 ng/mL) (Gibco, #200-10), GM-CSF (50 ng/mL) (Gibco, #300-03).

### Flow cytometry

#### Monocyte isolation profiling

Isolated monocytes (described in **Human primary cells**) were thawed and immediately washed with PBS. Cells were then stained using LIVE/DEAD Aqua (ThermoFisher #L34966) for 30 minutes at room temperature. Cells were then washed twice, and Fc receptors were blocked using Fc Receptor Block (BioLegend, #422301) at a 1:20 dilution in FACS buffer (3% FBS and 5mM EDTA in PBS) on ice for 20 minutes. Staining was performed in the Fc Binding Inhibitor + FACS buffer solution for 1 hour at 4°C using the following pre-conjugated monoclonal antibodies: CD45-eFluor450, clone HI30, 1:200 (eBioscience), CD16-PE, clone CB16, 1:200 (eBioscience), and CD14-APC, clone 61D3, 1:200 (eBioscience). After staining, cells were washed twice with FACS buffer and analyzed using an Attune NxT Flow Cytometer (Blue/Red/Yellow/Violet 6) (Thermo Fisher). FlowJo software (BD) was used for data analysis.

#### Macrophage surface marker profiling

Macrophages were lifted with Accutase (STEMCELL Technologies #07920) for 30 minutes at 37°C. Cells were then washed with PBS and stained using LIVE/DEAD Near-IR (ThermoFisher #L34975) for 30 minutes at room temperature. Cells were then washed twice, and Fc receptors were blocked using Fc Receptor Block (BioLegend, #422301) at a 1:20 dilution in FACS buffer (3% FBS and 5mM EDTA in PBS) on ice for 20 minutes. Staining was performed in the Fc Binding Inhibitor + FACS buffer solution for 1 hour at 4C using the following pre-conjugated monoclonal antibodies: CD80-PE, clone 2D10.4, 1:200 (eBioscience), CD206-BV421, clone 19.2, 1:200 (eBioscience), CD163-APC, clone GHI/61, 1:200 (eBioscience), CD40-PE-Cy7, clone 5C3, 1:200 (eBioscience), and PDL1-PerCP-eFluor710, clone MIH1, 1:200 (eBioscience). After staining, cells were washed twice with FACS buffer and analyzed using an Attune NxT Flow Cytometer (Blue/Red/Yellow/Violet 6) (Thermo Fisher). FlowJo software (BD) was used for compensation and gating. For analysis, samples with low transduction efficiency (“% GFP^+^ living cells” < 62.5%) were filtered out.

### Cell lines

K562, BT474, and SKOV3 cell lines were originally commercially obtained from the American Type Culture Collection (ATCC). Lx293T cells were purchased from Takara Bio USA (#632180) and cultured in DMEM (Life Technologies #10564029) + 10% FBS (Gibco, #A3840102) + 50μg/mL Gentamycin (Corning #30-005-CR). Both K562 cells and BT474 cells were cultured in complete RPMI 1640 medium supplemented with GlutaMAX, HEPES, and Phenol Red (Gibco, #72400047) with 10% fetal bovine serum (FBS) (Gibco, #A3840102). SKOV3 cells were cultured in complete McCoy’s 5A medium (Gibco, #16600082) with 10% FBS (Gibco, #A3840102). For expanding cells in culture, all cells were kept between 30-80% confluence. Adherent cell cultures were split using TrypLE (Gibco, #12604013). Cells were frozen using the following solution: 80% complete media (listed above), 10% FBS (Gibco, #A3840102), 10% DMSO (Sigma-Aldrich, #D8418). All tissue culture was performed under sterile conditions. All cells were kept in temperature-controlled incubators at 37°C with 5% CO_2_.

To generate BFP^+^ K562, BT474, and SKOV3 cells, each cell line was transduced with lenti-NLS-mTagBFP2-P2A-ffluc (described in **Lentiviral vector construction**). Cells were then sorted for high expression of BFP using fluorescence-activated cell sorting (FACS) at the Stanford Shared FACS Facility. BFP^+^/HER2^+^ K562 cells were provided by the lab of Dr. Rogelio Hernandez-Lopez. To generate HER2-overexpressing K562s, K562 cells were transduced with a lentiviral vector encoding both cDNA for HER2 and BFP. K562s with high expression of BFP were sorted using Fluorescence-activated cell sorting (FACS) at the Stanford Shared FACS Facility. For all cell lines expressing HER2, HER2 expression was validated using flow cytometry.

### *E. coli* phagocytosis assay

pHrodo-red *E. coli* Bioparticles (Invitrogen, #P35361) were thawed, resuspended, and added to macrophages at 100μg/mL in serum-free macrophage media (STEMCELL Technologies, #10961). Cells were promptly imaged using a 4X objective on either the Cellcyte (Echo, CellCyte 1) or the Incucyte (Sartorius, Incucyte S3), every 30-45 minutes for 24 hours. Red fluorescence intensity measurements were downloaded from the respective imager using either CELLCYTE studio software (v.2.7.4) or Incucyte S3 software (v2018B). “*E. coli* phagocytosis index” was calculated using either “red total intensity (a.u)” (CellCyte) or “red total integrated intensity (RCU x µm²/Image)” (Incucyte). For analysis, samples with low transduction efficiency (“green count per image” < 1000) were filtered out.

### Bulk RNA sequencing of primary macrophages

Monocytes from three different donors were isolated, differentiated, and transduced with either pHR-PGK-SCR-T2A-GFP, pHR-PGK-SCR(S3-S3)-T2A-GFP, or pHR-PGK-SCR(S1-S1)-T2A-GFP, and the media was changed the following day (described in **Human primary cells** and **Primary macrophage transduction**). Cells were then incubated for 3 days, after which they were treated with either IL-10 (50 ng/mL) (Gibco, #200-10) or IFN-γ (50 ng/mL) (STEMCELL Technologies, #78020). Cells receiving cytokine treatment were transduced with the control pHR-PGK-SCR-T2A-GFP to ensure transcriptional changes between samples were not due to lentiviral transduction. 48 hours after cytokine treatment, cells were imaged on the Cellcyte (Echo, CellCyte 1) to verify high (>80%) and even transduction efficiency across samples. Total RNA was extracted from each well using the RNeasy Mini Kit (QIAGEN, #74104). cDNA library construction using mRNA library preparation (poly A enrichment) and bulk RNA sequencing were performed by Novogene (Sacramento, CA) on an Illumina NovaSeq X Plus System (PE150). MultiQC^48^ was used to evaluate sequencing quality with default parameters. Transcript abundances were quantified using Salmon (v1.10.3)^49^, a reference transcriptome (Gencode GRCh38.p14), and default parameters. Differential gene expression analysis (DESeq2)^50^, Gene Set Enrichment Analysis (GSEA)^51^, Kyoto Encyclopedia of Genes and Genomes (KEGG)^52^, and Gene Ontology (GO)^53,54^ pathway enrichment analysis were performed in R (v4.3.2). MSigDB (v3.0)^55^ was used to identify lists of genes experimentally validated to be regulated by IL-10 and IFNγ. Hierarchical clustering was performed using the “pheatmap” function in R. Transcription factor activity was inferred from bulk RNA sequencing data using the DoRothEA human regulon database^56^ and the decoupleR weighted mean method^57^. TF activity scores were calculated for each sample and contrasted using linear modeling (limma) relative to the reference condition SCR0. Empirical Bayes moderation was applied, and p-values were adjusted using the Benjamini–Hochberg method.

#### Gene signature scoring and cytokine analysis from scRNA-seq data

Publicly available scRNA-seq (GSE186802) from diffuse midline glioma patients treated with GD2-CAR T therapy^34^ was downloaded and analyzed in R using Seurat (v5)^58^. Data were normalized (‘LogNormalize’, scale factor 10,000), scaled (‘ScaleData’), and UMAPs generated. The CAR T-cell product was removed from the dataset based on metadata. Myeloid cells were specifically selected based on relatively high expression of myeloid genes (AIF1, CSF1R, CD14, CD68), compared to B cell-related genes (CD19, CD22, PAX5), and T cell-related genes (CD2, CD3E, CD3D, CD3G, CD247, and CD7). Differential gene expression analysis (DESeq2)^50^ was used to create a signature of genes upregulated in SCR(S1-S1) vs SCR0 (p.adj <0.05, baseMean > 50, log2FoldChange >=2). Gene signatures were analyzed using ‘AddModuleScore_Ucell’ and hierarchical clustering was performed using “pheatmap”. Donor (i.e., DIPG1, DIPG2, DIPG3, and spinal_DMG1) and treatment (intravenous (IV) or intracerebroventricular (ICV) infusion) information was stated in the original publication.

IFNγ measurements from the cerebrospinal fluid of the same were extracted from source data^34^ and matched to scRNA-seq sampling timepoints. Cytokine values were normalized to day 0 levels for each patient and each infusion type and expressed as log2 fold-change before downstream correlation analyses.

### CAR macrophage and cancer cell co-culture assays

For the BT474 co-culture assay, BFP^+^ BT474 cells were generated as described in **cell lines**. Macrophages (50,000 cells per well) were cultured in a Nunc dish with an UpCell surface (Thermo Scientific). Four days after macrophages were transduced as described in **primary macrophage transduction > in vitro experiments**, BT474 cells were pH-labeled using pHrodo Red SE (Invitrogen, #P36600). The pHrodo-labeled BT474s were added at a specified effector to target (E:T) ratio using 1/2 media indicated for the cancer cell line of interest and 1/2 serum-free macrophage media (STEMCELL Technologies #10961) by volume, with M-CSF (STEMCELL Technologies #78057) at 50ngs/mL. For the first two challenges, 25,000 BT474 cells (E:T = 2:1) were added to each well. For the second two challenges, 50,000 BT474 cells (E:T = 1:1) were added to each well. Wells were imaged (BF, Blue, Red, and Green channels) using a 4X objective on the Cellcyte 1 (Echo), every 3 hours for 12 days.

For the K562 co-culture assay, HER2^+^ K562s were generated as described in **cell lines**. Macrophages (50,000 cells per well) were cultured in a Nunc dish with an UpCell surface (Thermo Scientific). Four days after macrophages were transduced as described in **primary macrophage transduction > in vitro experiments**, HER2^-^ or HER2^+^ K562s (20,000 per well) were added to each well. Wells were imaged (BF, Blue, and Green channels) using a 4X objective on the Cellcyte 1 (Echo), every 3 hours for 3 days.

### Animal models

All experimental studies were approved and performed in ethical compliance with Institutional Animal Care and Use Committee (IACUC) approved protocols at Stanford University. NOD/SCID/IL2Rγ^−/−^ (NSG) mice were purchased from JAX and bred by the Breeding Colony Management Service at Stanford University. Mice were housed in sterile cages at 22°C and 50% humidity under a 12h-12h light-dark cycle. Mice were monitored daily by the Veterinary Services Center (VSC) staff at Stanford University. Mice that were used for experiments were not involved in previous procedures or given drug treatment. Mice were aged between 6 and 10 weeks at tumor or macrophage engraftment. Mice were euthanized upon manifestation of persistent hunched posture, scruffy coat, paralysis, impaired mobility, or greater than 20% weight loss. Sample size calculations were not performed, but were determined by experience with well-established, previously published models^36,59^.

### SKOV3 ovarian cancer model

1 x 10^6^ SKOV3 cells in 100μL of PBS were injected intra-peritoneally (IP) into female mice aged 8-10 weeks old. Tumor growth was monitored using bioluminescent imaging (BLI), with the first image taken between SKOV3 injection and macrophage injection. Mice were then split into random groups of five mice each and injected with 100uL of PBS or 3 x 10^6^ macrophages in PBS (preparation of macrophages is described in **Primary macrophage transduction > In vivo experiments**). Tumor growth was monitored using BLI.

### Bioluminescent imaging

Mice were injected with 50μL of 15mg mL^-1^ D-Luciferin retro-orbitally to avoid interfering with cancer cells or macrophages in the peritoneal cavity. Images were acquired on the IVIS (Perkin Elmer) imaging system immediately after injection using exposure times of 1 second, 15 seconds, 30 seconds, and auto-exposure settings. Total flux was measured using Living Image (v4.8.2; Perkin Elmer) software by drawing a region of interest around the body of each mouse. Only non-saturated images were used for quantification of BLI. For visualization of tumor burden, images were set to the same luminescent scale.

### Luminex cytokine measurements

At the end of the third challenge of macrophages with BT474 cells, supernatant was collected for further processing. The Luminex assay was performed by the Human Immune Monitoring Center at Stanford University - Immunoassay Team. Kits were purchased from EMD Millipore Corporation, Burlington, MA, and run according to the manufacturer’s recommendations with modifications described as follows: The H48 kits include one panel: Milliplex HCYTA-60K-PX48. The assay setup adhered to the recommended protocol with the following steps: 1. Sample Preparation: Supernatant samples were run undiluted. 2. Incubation: The samples were incubated overnight at 4°C with shaking. Cold and room temperature incubation steps were performed on an orbital shaker at 500-600 rpm. 3. Washing: Plates were washed twice with wash buffer using a BioTek ELx405 washer (BioTek Instruments, Winooski, VT). 4. Detection: After a one-hour incubation at room temperature with a biotinylated detection antibody, streptavidin-PE was added and incubated for 30 minutes with shaking. 5. Final Wash and Reading: Plates were washed as described above, and Wash Buffer was added to the wells for reading in the Luminex FlexMap3D Instrument, ensuring a lower bound of 50 beads per sample per cytokine. 6. Replication and Quality Control: Each sample was measured in duplicate replicates. Custom Assay Chex control beads (Radix BioSolutions, Georgetown, Texas) were added to all wells. Wells with a bead count <20 were excluded.

### Two-state model

The two-state model was carried out using both R (v.4.3.2) and Mathematica (Wolfram), which produced nearly identical results. Motif combinations and all phenotype measurements (CD163, CD206, CD80, CD40, PDL1, and phagocytosis) for all replicates were used as input to fit models that predict the effect of SCR motif combinations on each phenotype. Equations used in this model can be found in **Extended Data Figure 7** and on Github (see **Code Availability**). For each SCR, the copy number of each of the 9 motifs (S1–S9) was enumerated from the motif composition metadata. Standard errors across technical replicates were retained as a measure of replicate variability but were not used as weights in fitting. Phenotype-specific fluorescence bounds were defined as the single lowest and single highest mean observed value across all samples for that phenotype, specifying the dynamic range of the two-state model. Model parameters—an intrinsic energy term (ΔEi) and a per-motif energy contribution for each of the nine motifs (ΔEs for S1–S9)—were estimated by minimizing the sum of squared residuals between observed and model-predicted fluorescence values, using all individual technical replicates and BFGS optimization (`optim()` in base R). Model fit statistics (r and R²) were evaluated on per-condition averages of technical replicates. Generalization performance was evaluated via 10-fold cross-validation using the `rsample` package. Coefficient stability was assessed as the standard deviation of each ΔEs estimate across folds. The difference between in-sample R² and cross-validated R² was used as a diagnostic for overfitting. Data manipulation was performed using `dplyr`, `tidyr`, and `purrr`. Per-motif equilibrium constant contributions (Ks) were derived from the fitted ΔEs coefficients, where Ks values greater than 1 indicate that a motif shifts the equilibrium toward the ON state, while values less than 1 indicate a shift toward the OFF state. Ks values are reported on a log₁₀ scale for visualization. Uncertainty on each log₁₀ Ks value was propagated from the cross-validation fold standard deviation of ΔEs. See **Data Availability** and **Code Availability** for details on how this model can be run.

### Statistical analysis

For all experiments, the number of replicates and statistical analyses used are described in the figure legends. Statistical analyses were performed using R (v.4.3.2). P value <0.05 was considered statistically significant (p < 0.05, ** p < 0.01, *** p < 0.001).

## Data Visualization

Initial plots were generated using R (v.4.3.2). Plots were organized and refined with Adobe Illustrator (v.30.2.1).

## Data Availability

Source data are provided with this paper as of the date of publication. The data for the bulk RNA-seq dataset described in this study have been deposited at the NCBI Gene Expression Omnibus (GEO) under accession number GSE328683. Differentially expressed genes from bulk RNA sequencing, as well as surface marker and phagocytosis measurements which can be used to run the two-state model are available at Zenodo (10.5281/zenodo.19699424) and will be publicly released upon publication.

## Code Availability

The custom code used to run the two-state model in this manuscript is available at https://github.com/jodielunger/SCR-macrophages and will be archived on Zenodo (10.5281/zenodo.19699424) upon publication.

## Acknowledgements

We thank Trung Pham and Karla Kirkegaard for insightful discussions. We thank the Barsh, Baker, Kirkegaard, Sherlock, and Hernandez-Lopez labs for generous sharing of equipment and reagents. We thank Jaclyn Ng and Céline Prange for offering support for mouse experiments and providing protocols. We thank Jaclyn Ng, Céline Prange, Adrià Cañellas-Socias, Kylie Burdsall, and Peng Xu for guidance with animal experiments. This work was supported by the grants listed below.

National Institutes of Health Grant DP2EB039061 (K.G.D)

National Institutes of Health Grant R21EB037367 (K.G.D)

Hypothesis Fund (K.G.D)

Shurl and Kay Curci Foundation (K.G.D)

Parker Institute for Cancer Immunotherapy (K.G.D)

Stanford CShaRP (K.G.D)

Stanford Cancer Institute (K.G.D)

Stanford Department of Genetics (K.G.D)

National Human Genome Research Institute 5T32HG000044-27 (J.C.L)

National Science Foundation Graduate Research Fellowship DGE-2146755 (A.S.A, L.E.S)

Stanford Graduate Fellowship (A.S.A)

## Author contributions

K.G.D and J.C.L conceptualized the study. J.C.L conducted experiments. J.C.L and K.G.D set up computational pipelines and analyzed all data. L.E.S and A.S.A provided support for the design and execution of *in vitro* experiments. R.A.H developed analysis to integrate the clinical scRNA-seq data with the bulk RNA-seq. J.Y.L cloned and tested initial designs of constructs used in this study. J.P.G cloned several constructs used in this study and provided support for *in vivo* experiments. A.V.J, M.G, W.C, and A.N.B provided support for the study and experimental design. All figures were prepared by J.C.L. The manuscript was written by J.C.L and K.G.D. The manuscript was finalized with input from all authors.

## Competing interests

K.G.D and J.C.L are co-inventors on a patent application related to the work described in this manuscript. No other authors declare competing interests.

---

**Extended Data Figure 1.**
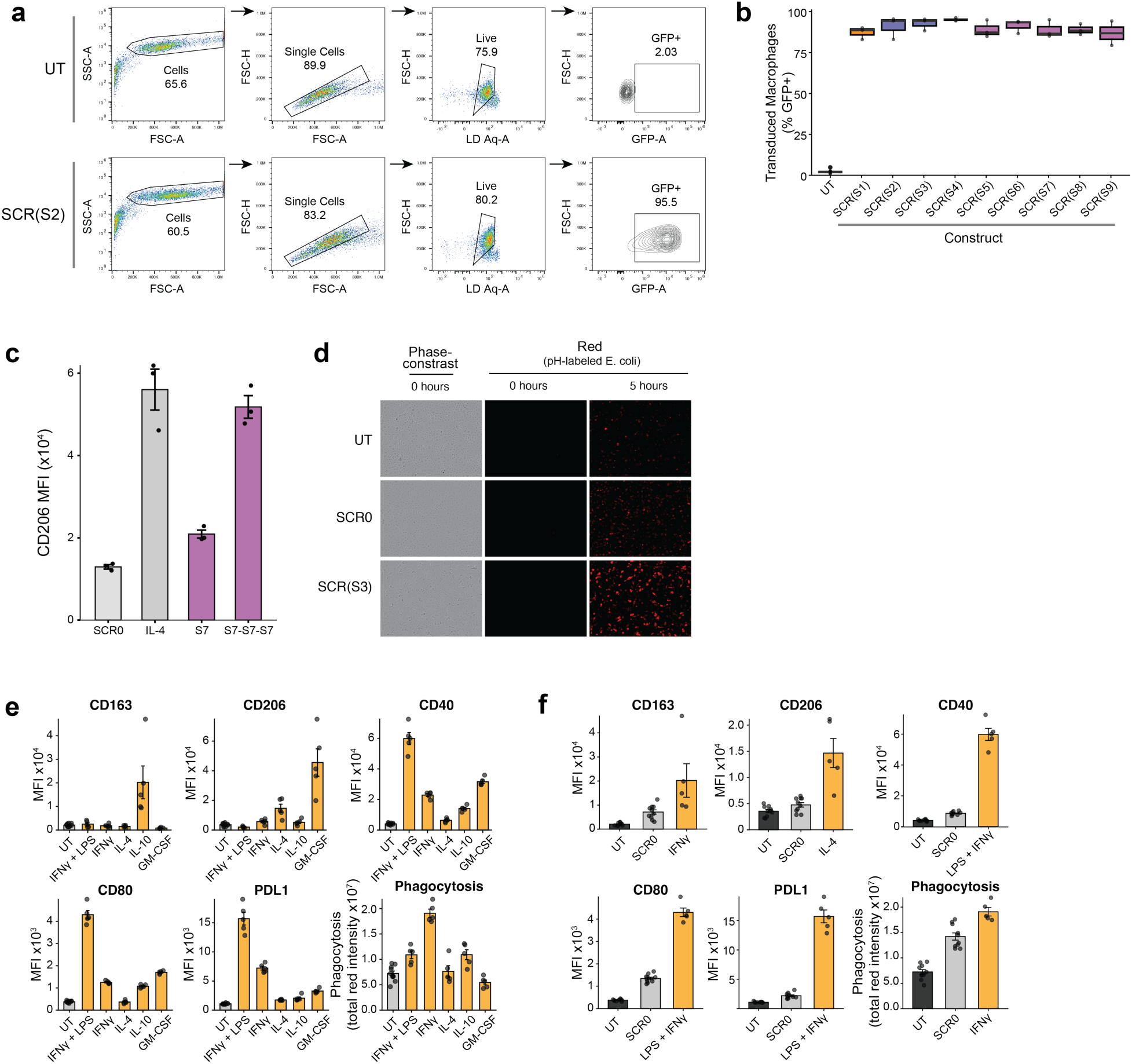
| Characterization of exogenous ligand-treated and SCR-expressing macrophages. **a,** Representative flow cytometry gating strategy for either an untransduced macrophage sample (UT) or macrophages transduced with SCR(S2). **b,** Transduction efficiency of each single-motif SCR as measured by the percentage of cells positive for GFP. **c,** MFI of CD206 for IL-4-treated macrophages compared to macrophages expressing an SCR containing no motif (SCR0), motif S7 in one or three copies. Data are mean ± s.e.m. MFI, mean fluorescence intensity. **d,** Representative images of phagocytosis of pHrodo-Red-labelled *E. coli* by untransduced macrophages (UT), macrophages transduced with SCR0, or macrophages transduced with SCR(S3). The initial (0 hours) timepoint was taken immediately after the addition of phRodo-Red-labeled *E.* coli. **e,** Quantification of MFI or phagocytosis of pHrodo-Red-labelled *E. coli* by untransduced macrophages (UT) or macrophages treated for 48 hours with the indicated exogenous ligands (orange). Data are mean ± s.e.m. MFI, mean fluorescence intensity. **f,** Quantification of MFI or phagocytosis of pHrodo-Red-labelled *E. coli* by untransduced macrophages (UT, black), macrophages transduced with an SCR containing no signaling motifs (SCR0, grey), or macrophages treated for 48 hours with the indicated exogenous ligands (orange). Data are mean ± s.e.m. MFI, mean fluorescence intensity.

**Extended Data Figure 2.**
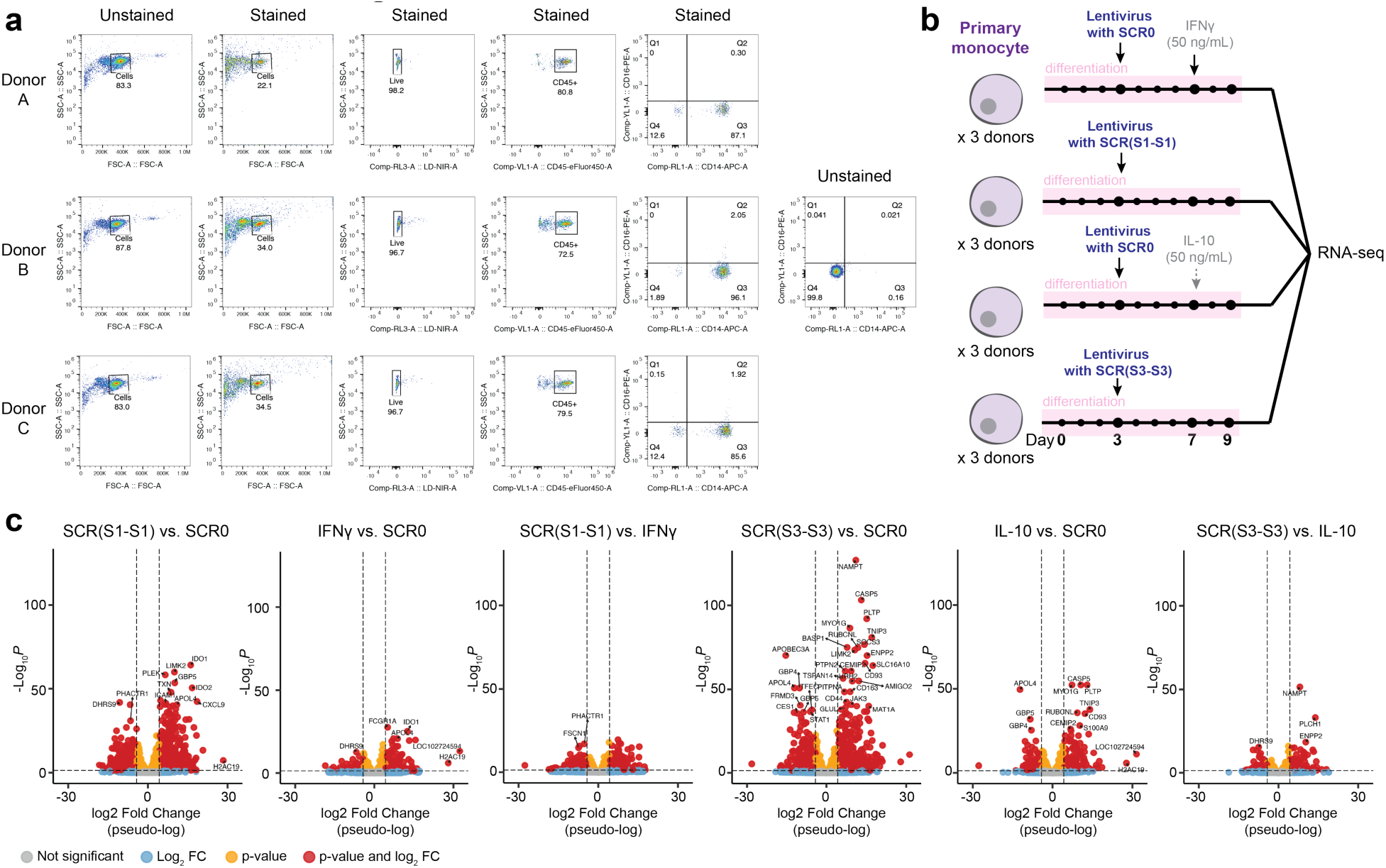
| RNA-sequencing comparing SCR-driven vs. cytokine-mediated transcriptional states. **a,** Representative flow cytometry plots for all three donor monocyte isolations used for the RNA-seq experiment. Cells were stained with CD45, CD14, and CD16. Unstained cells are indicated as controls. **b,** Schematic of experimental design for RNA-seq. RNA-seq, RNA sequencing. **c,** Volcano plots showing differential gene expression for each indicated comparison. Each point represents a single gene, plotted by log₂ fold change and −log₁₀ Benjamini–Hochberg–adjusted *p*-value. Genes meeting significance thresholds (adjusted p < 0.05 and log₂ fold change > 1) are highlighted in color. Selected, highly differentially expressed genes are labeled. Horizontal and vertical dashed lines indicate significance and fold-change cutoffs, respectively.

**Extended Data Figure 3.**
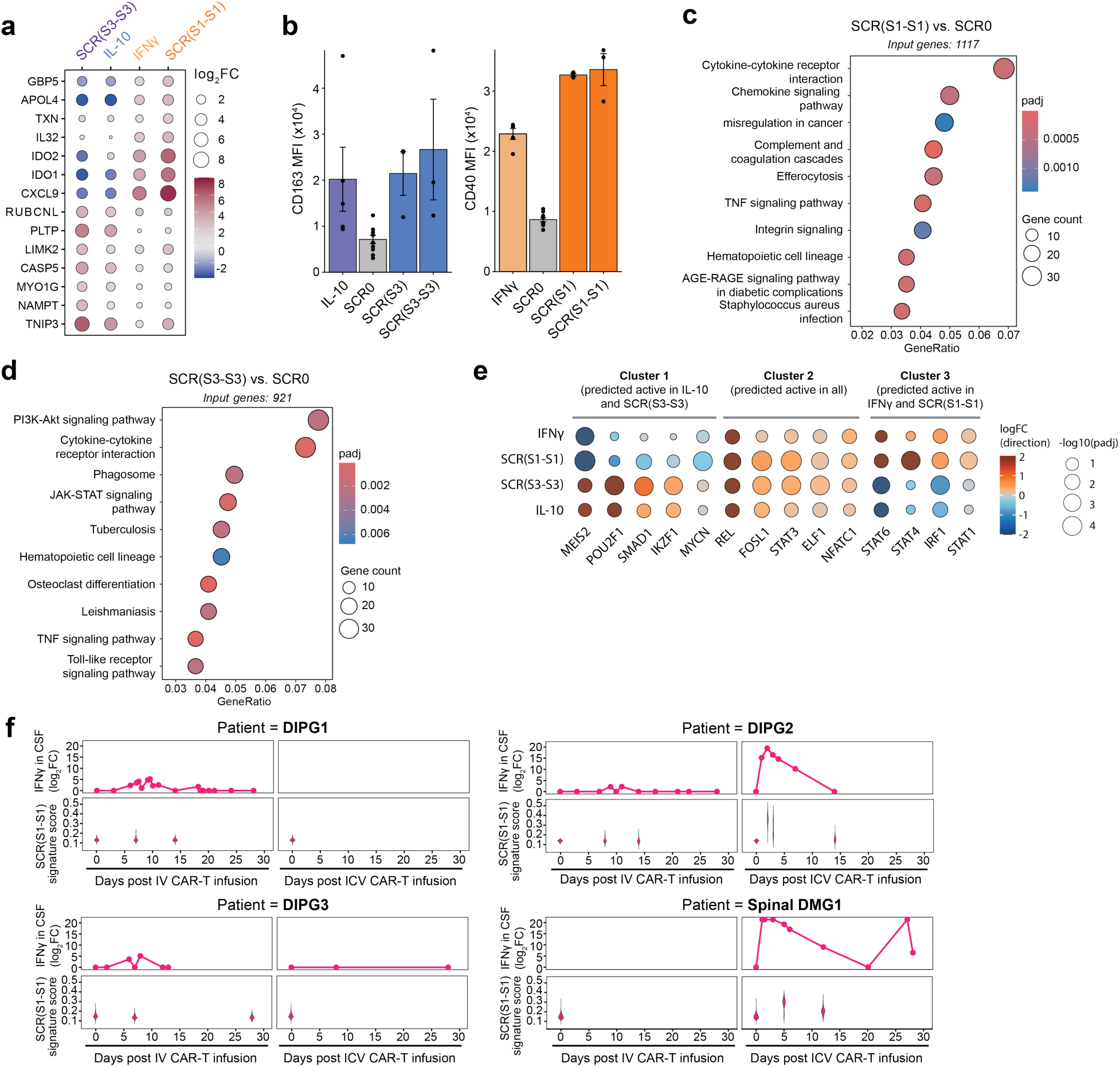
| Gene programs are differentially enriched in SCR(S1-S1) and SCR(S3-S3) conditions. **a,** MFI of CD163 for IL-10-treated macrophages compared to macrophages expressing an SCR containing no motif (SCR0), motif S3 in one (SCR(S3) or two (SCR(S3-S3) copies (left), or of CD40 for IFN-γ-treated macrophages compared to macrophages expressing an SCR containing no motif (SCR0), motif S1 in one (SCR(S1) or two (SCR(S1-S1) copies (right) as measured by flow cytometry. Data are mean ± s.e.m. MFI, mean fluorescence intensity. **b,** Bubble plot of the top 5 differentially expressed genes between each sample and the SCR0-expressing control (ranked by Benjamini–Hochberg–adjusted *p*-value). The color and size of the dot corresponds to the log_2_-fold-change (log_2_FC) of the indicated gene. FC, fold change. **c,** KEGG pathway enrichment analysis on differentially expressed genes identified for SCR(S1-S) relative to SCR0 control. The top 10 KEGG-enriched pathways are shown. Dot color represents Benjamini–Hochberg–adjusted p-value (padj). The total number of genes used as input for analysis is indicated. **d,** KEGG pathway enrichment analysis on differentially expressed genes identified for SCR(S1-S) relative to SCR0 control. The top 10 KEGG-enriched pathways are shown. Dot color represents Benjamini–Hochberg–adjusted p-value (padj). The total number of genes used as input for analysis is indicated. **e,** Bubble plot of inferred transcriptional activity from bulk RNA sequencing using DoRothEA human regulons. For each contrast (indicated condition vs. mock transduction control (SCR0), the top five transcription factors (lowest padj) were selected, and the union of these transcription factors across all contrasts is shown. Bubble size represents statistical significance (−log10 padj). Color represents the direction and magnitude of activity change. FC, fold change. **f,** Fold change of IFNγ in the CSF relative to day 0, after either IV or ICV CAR T infusion (above), compared to the SCR(S1-S1) signature of myeloid cells in the CSF for each of the four patients reported in Majzner and Ramakrishna et al, 2022^34^. IV, intravenous; ICV, intracerebroventricular; FC, fold change; DIPG, diffuse intrinsic pontine glioma; DMG, diffuse midline glioma; CSF, cerebrospinal fluid.

**Extended Data Figure 4.**
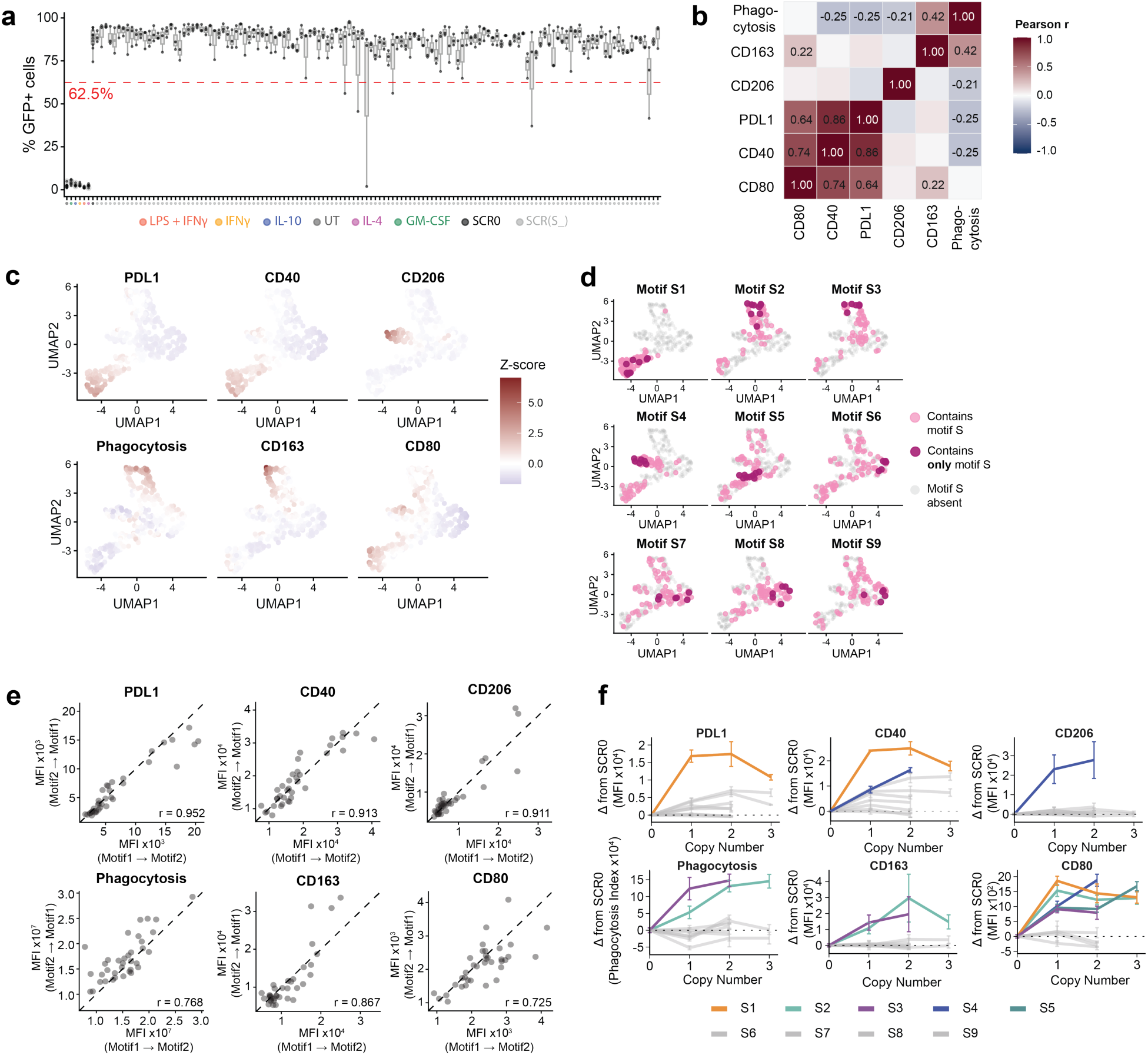
| Combining motifs on the SCR reveals a navigable landscape of primary macrophage polarization states. **a,** Transduction efficiency across all samples, measured by the percentage of living macrophages that are positive for GFP (% GFP^+^ cells). The red dashed line indicates the threshold of 62.5% applied to the dataset for further analysis. **b,** Heatmap of the pairwise Pearson correlation coefficients between all measured phenotypes. Color intensity indicates the strength and direction of the correlation. Select correlation coefficients are displayed within each cell. **c,** UMAP from Fig. 3e containing only points for macrophages expressing an SCR from the combinatorial library, colored by z-score for all measured phenotypes. **d,** UMAP projection colored by motif presence for all motifs. Pink indicates SCRs containing the indicated motif, while maroon indicates SCRs containing only that motif in one, two, or three copies. S, signaling motif. **e,** Pairwise correlations for all double-motif combinations and all phenotypes measured. Pearson correlation coefficient (r) is indicated on each plot. MFI, mean fluorescence intensity. **f,** Line plots showing the effect of increasing copy number (0–3) of each signaling motif on all measured phenotypes. Relative change from SCR0 (Δ) was calculated as the difference between the mean phenotype value for each motif copy number and the mean value of the SCR0 control. Colors indicate the signaling motif. Data are mean ± s.e.m. MFI, mean fluorescence intensity.

**Extended Data Figure 5.**
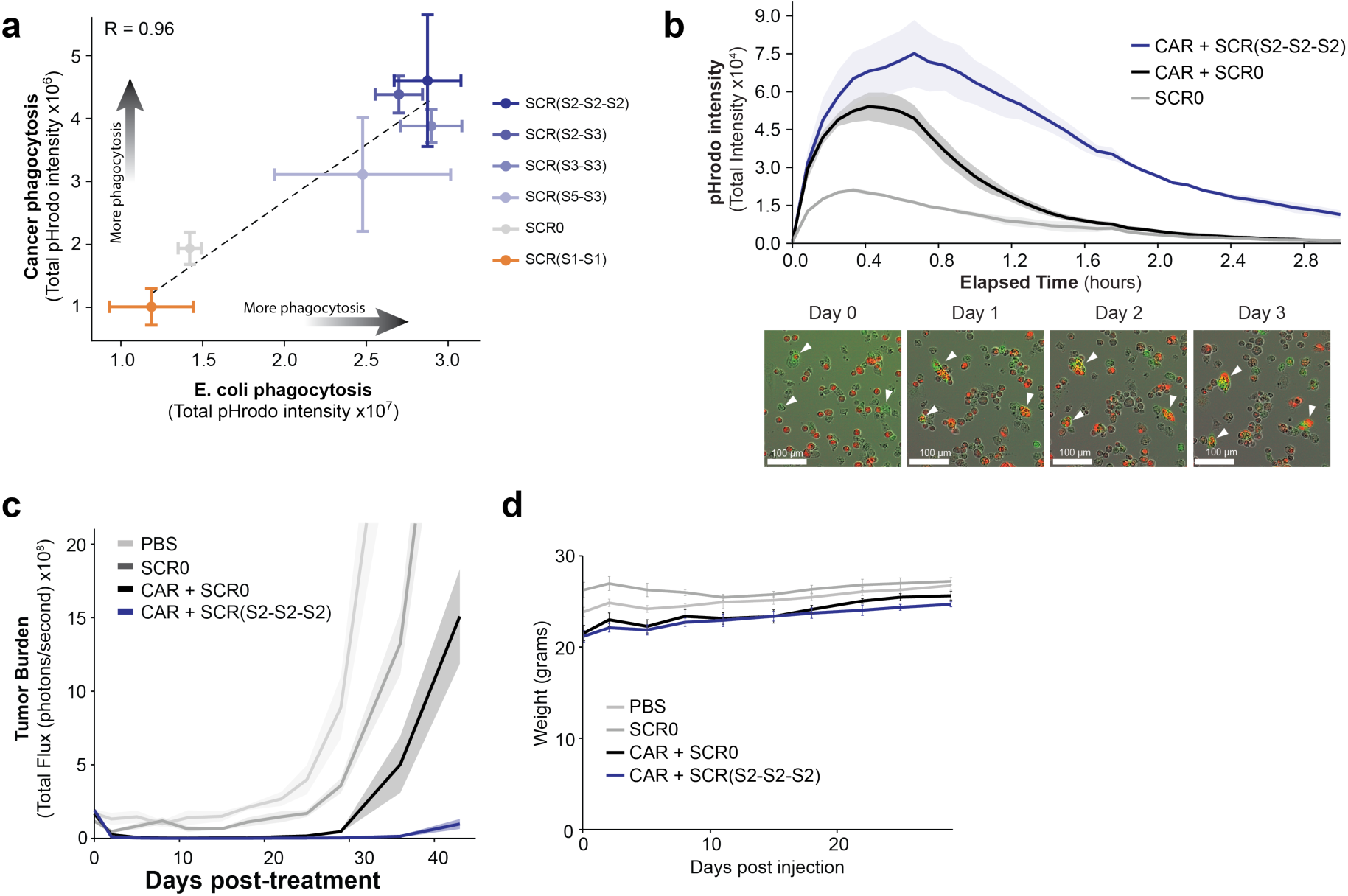
| SCR expression augments phagocytosis of cancer cells by CAR-Ms. **a,** Correlation between phagocytosis of *E. coli* in Fig. 3d and phagocytosis of cancer by CAR-Ms in Fig. 4f**. b,** Quantification of pHrodo intensity and representative images from single-challenge co-culture of pHrodo-labeled HER2^+^ K562s and macrophages transduced with SCR0 (grey, *n* = 2), CAR + SCR0 (black, *n* = 2), or CAR + SCR(S2-S2-S2) (blue, *n* = 2). Data are mean ± s.e.m. **c,** Tumor burden as seen in Fig. 4j on a linear scale. Tumor burden is measured as the total flux (photons s⁻¹) of luciferase signal. Data are mean ± s.e.m. **d,** Weights of mice over the course of the *in vivo* experiment depicted in Fig. 4h. Data are mean ± s.e.m.

**Extended Data Figure 6.**
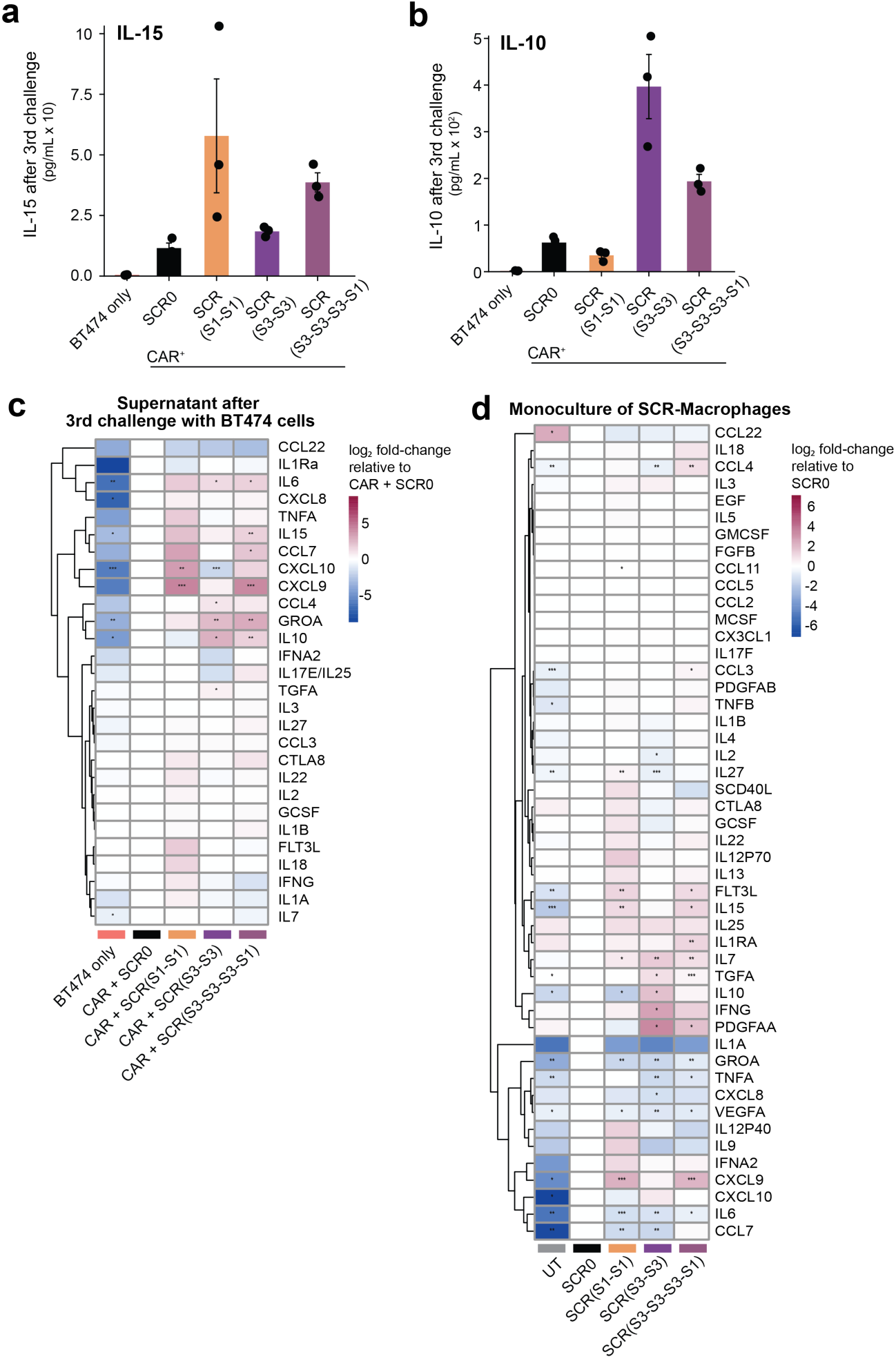
| Model-guided SCR drives a unique cytokine secretion profile. **a,** Concentration of IL-15 in the supernatant at the end of the third addition of BT474 cells to CAR macrophages co-expressing the indicated SCR. Data are mean ± s.e.m. **b,** Concentration of IL-10 in the supernatant at the end of the third addition of BT474 cells to CAR macrophages co-expressing the indicated SCR. Data are mean ± s.e.m. **c,** Heatmap showing log_2_ fold-change in cytokine concentration in the supernatant at the end of the third addition of BT474 cells to CAR macrophages co-expressing the indicated SCR, relative to the CAR + SCR0 control. Only cytokines with minimal baseline secretion by BT474 cells alone are shown. Statistical significance was assessed using a two-sample t-test comparing replicate concentration values (n = 3) for each condition against the CAR + SCR0 baseline for each cytokine independently. **d,** Heatmap showing log_2_ fold-change in cytokine concentration in the supernatant of macrophage monoculture, four days after transduction with the indicated SCR. Statistical significance was assessed using a two-sample t-test comparing replicate concentration values (n = 3) for each condition against the SCR0 baseline for each cytokine independently.

**Extended Data Figure 7.**
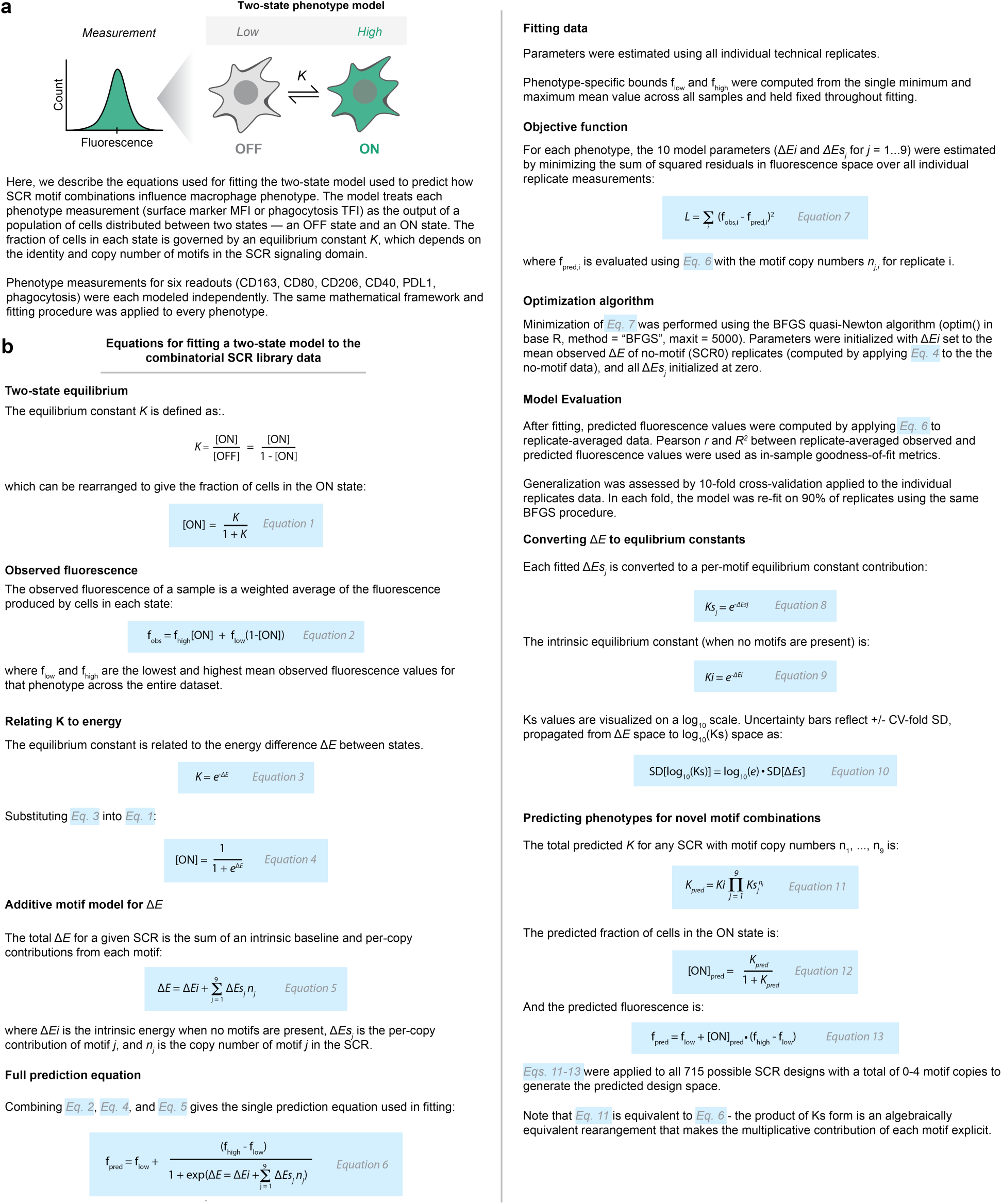
| A two-state model for predicting motif-to-phenotype relationship. **a,** Graphic and high-level explanation of the two-state model framework. **b,** Equations used for fitting parameters and model evaluation.

## Notes

### Summary of Updates

Some typos have been fixed. Some small edits to the figures have been made for clarity.

